# Deep mutational scanning reveals characteristics important for targeting of the tail-anchored protein Fis1

**DOI:** 10.1101/045351

**Authors:** Abdurrahman Keskin, Emel Akdoğan, Cory D. Dunn

## Abstract

Proteins localized to mitochondria by a carboxyl-terminal tail anchor (TA) play roles in apoptosis, mitochondrial dynamics, and mitochondrial protein import. To reveal characteristics of TAs that may be important for mitochondrial targeting, we focused our attention upon the TA of the *Saccharomyces cerevisiae* Fis1 protein. Specifically, we generated a library of Fis1p TA variants fused to the Gal4 transcription factor, then, using next-generation sequencing, revealed which Fis1p TA mutations inhibited membrane insertion and allowed Gal4p activity in the nucleus. Prompted by our global analysis, we subsequently analyzed the ability of individual Fis1p TA mutants to localize to mitochondria. Our findings suggest that the membrane-associated domain of Fis1p TA may be bipartite in nature, and we encountered evidence that the positively charged patch at the carboxyl-terminus of Fis1p is required for both membrane insertion and organelle specificity. Furthermore, lengthening or shortening the Fis1 TA by up to three amino acids did not inhibit mitochondrial targeting, arguing against a model in which TA length directs insertion of TAs at specific organelles. Most importantly, positively charged residues were more acceptable at several positions within the membrane-associated domain of the Fis1p TA than negatively charged residues. These findings, emerging from the first high-resolution analysis of an organelle targeting sequence by deep mutational scanning, provide strong, *in vivo* evidence that lysine and arginine can “snorkel,” or become stably incorporated within a lipid bilayer by placing terminal charges of their side chains at the membrane interface.

**Abbreviations:**

TA: tail anchor
OM: outer membrane
MAD: membrane-anchoring domain
3-AT: 3-aminotriazole
CHX: cycloheximide

## INTRODUCTION

Proteins inserted within the mitochondrial outer membrane (OM) by a carboxyl-terminal tail anchor (TA) are important for programmed cell death, mitochondrial protein import, and the control of mitochondrial shape and number (Wattenberg and Lithgow 2001). While ER-directed tail-anchored proteins can take advantage of a conserved set of soluble proteins and membrane-bound receptors (Denic *et al.* 2013; Johnson *et al.* 2013), the machinery targeting many TAs to mitochondria is yet to be discovered (Lee *et al.* 2014; Neupert 2015), and genetic and biochemical evidence suggest that spontaneous insertion of TAs at mitochondria may occur without the need for a translocation machinery (Setoguchi *et al.* 2006; Kemper *et al.* 2008). Furthermore, some tail-anchored proteins can be dual-localized to mitochondria and other organelles (Borgese and Fasana 2011), but how membrane specificity is controlled is unclear. TA targeting seems to depend, in general, upon incompletely defined structural characteristics of the TA rather than a defined consensus sequence (Beilharz 2003; Rapaport 2003; Borgese *et al.* 2007).

Genetic selection schemes using the organism *Saccharomyces cerevisiae* have been of high value in understanding how proteins reach their proper destination within eukaryotic cells. During such studies, a protein required for survival under selective conditions can be mislocalized, and thereby made inactive, by a targeting sequence utilized by the transport process being studied. Next, mutations that allow return of this mistargeted protein to a region of the cell at which it can perform its function are recovered under selective conditions. *Trans* factors related to protein targeting are identified by standard genetic approaches. Alternatively, *cis* mutations in the targeting sequence are revealed, typically by Sanger sequencing of individual fusion construct clones. Most prominently, this genetic approach to studying protein targeting and transport has been important in understanding protein transit to and through the endomembrane system (Deshaies and Schekman 1987; Robinson *et al.* 1988; Stirling *et al.* 1992). This approach has also been applied to the study of protein import and export at the mitochondrial inner membrane (Jensen *et al.* 1992; Maarse *et al.* 1992; He and Fox 1999).

Even with the availability of these powerful genetic strategies, a fine-grained analysis of any single eukaryotic protein targeting signal has been lacking. However, with the advent of next-generation sequencing, more comprehensive studies of protein targeting sequences are possible. In this study, we successfully coupled genetic selection to next-generation sequence analysis in order to define characteristics important for localization of the tail-anchored Fis1 protein to the mitochondrial outer membrane (OM).

## RESULTS

### Localization to mitochondria via the Fis1 tail anchor prevents Gal4-mediated transcriptional activation

The TA of Fis1p is necessary (Mozdy *et al.* 2000; Beilharz 2003) and sufficient (Kemper *et al.* 2008; Förtsch *et al.* 2011) for insertion of this polypeptide into the mitochondrial OM. No cellular machinery involved in Fis1p insertion has been identified (Kemper *et al.* 2008). Fis1p has been suggested to reach a final topology in the outer membrane in which the amino-terminal bulk of the protein faces the cytosol, a very short and positively charged carboxyl-terminus protrudes into the mitochondrial intermembrane space, and the two are connected by a membrane-anchoring domain (MAD) passing through the OM (Mozdy *et al.* 2000). In developing our selection for TA mutations that diminish Fis1p targeting, we reasoned that fusion of a transcription factor to Fis1p would lead to insertion within the mitochondrial OM and a lack of nuclear function (Figure 1A). Mutations within the TA of Fis1p that prevent effective membrane insertion would, however, presumably allow the linked transcription factor to enter the nucleus, promote expression of its targets, and allow survival under specific selective conditions, provided that the fusion protein is not degraded, aggregated, or misdirected to another cellular location. Toward this goal, we generated a construct containing the Gal4 transcription factor at the amino terminal end of the polypeptide and full-length Fis1p at the carboxyl terminal end of the protein, since *S. cerevisiae* strains allowing titratable selection based upon nuclear entry and subsequent binding to Gal4p-responsive DNA elements are readily available. Superfolder GFP (Pédelacq *et al.* 2005) was placed between the Gal4 and Fis1 moieties and was visible at mitochondria upon overexpression of this fusion protein (Figure 1B). While Fis1p has been reported to be homogenously distributed on the mitochondrial surface (Mozdy *et al.* 2000), puncta containing Gal4-sfGFP-Fis1p are observed, perhaps due to the formation of heteromeric complexes with nuclear import components attempting to transport Gal4-sfGFP-Fis1p to the nucleus.

**Figure 1.**
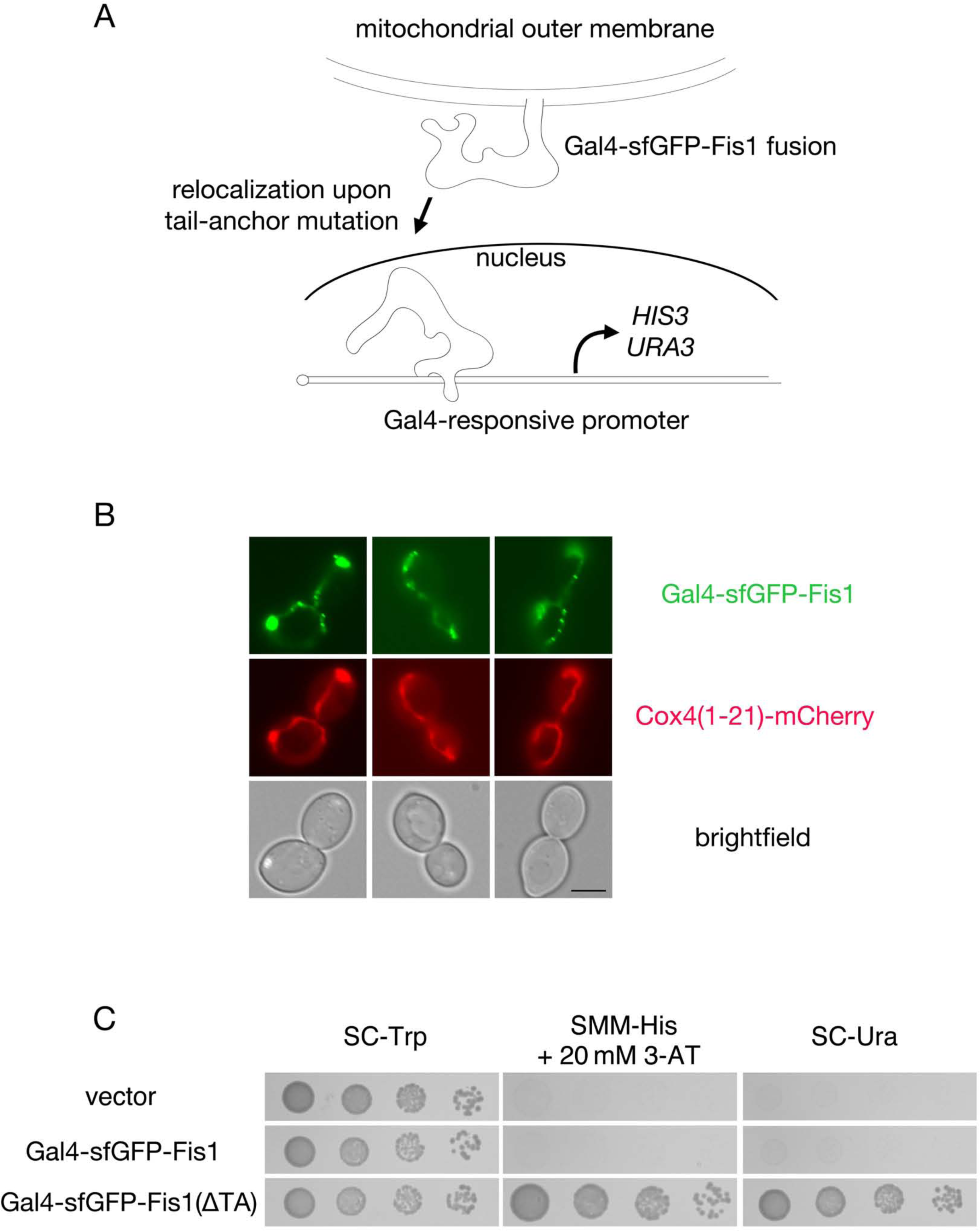
A genetic selection based on protein mislocalization allows recovery of mutations blocking Fis1p TA localization to mitochondria. (A) A scheme for selection of mutations preventing mitochondrial targeting of the Fis1p TA. Full-length Fis1p is fused to the transcription factor Gal4p. Upon failure of Fis1p to be localized to the mitochondrial OM, Gal4p may be free to translocate to the nucleus and activate the selectable markers *HIS3* and *URA3*. (B) A Gal4-sfGFP-Fis1 fusion localizes to mitochondria. *S*train CDD898 was transformed with plasmid b102, which overexpresses the Gal4-sfGFP-Fis1p construct used in this study. Mitochondria were visualized using mCherry fused to the Cox4 presequence expressed from plasmid pHS12-mCherry. Scale bar, 5 μm. (C) Removal of the Fis1p TA allows proliferation on medium requiring *HIS3* activation or *URA3* activation. Strain MaV203 expressing Gal4-sfGFP-Fis1p from plasmid b100 (WT TA), a variant lacking the Fis1p TA from plasmid b101 (ΔTA), or harboring empty vector pKS1 was cultured in SC-Trp medium, then, following serial dilution, spotted to SC-Trp, SMM-His + 20 mM 3-AT, or SC-Ura and incubated for 2 d.

To assess failed Gal4-sfGFP-Fis1p targeting to mitochondria, we specifically took advantage of the Gal4-driven *HIS3* and *URA3* auxotrophic markers in MaV203, a strain commonly used for yeast-two-hybrid assays (Vidal *et al.* 1996a). Similar to cells containing an empty vector, Gal4p fused to Fis1p was unable to allow proliferation on medium lacking histidine and containing 20mM 3-aminotriazole (3-AT) to competitively inhibit any His3p produced independently of Gal4p activation (Durfee *et al.* 1993) or in medium lacking uracil (SC-Ura) (Figure 1C). However, the same Gal4-sfGFP-Fis1 polypeptide devoid of its TA [Gal4-sfGFP-Fis1(ΔTA)] permitted ample proliferation on the same two selective media. This result indicated that our fusion protein could translocate to the nucleus upon TA disruption and that any potential lipid binding mediated by the cytosolic domain of Fis1p (Wells and Hill 2011) will not prevent genetic assessment of TA localization.

We immediately sought mutations in the TA that would block targeting of Gal4-sfGFP-Fis1p to the mitochondrial OM by isolating colonies spontaneously arising on medium lacking uracil. Limited sequencing of the constructs encoding Gal4-sfGFP-Fis1p within these isolates revealed at least seven nonsense and 19 frameshift mutations out of a total of 32 plasmids analyzed. While these findings further validated the link between proliferation under selective conditions and damage to the Fis1p TA, continued encounters with nonsense and frameshift mutations would not be greatly informative regarding sequential or structural determinants important for TA targeting. As expected, we noted that release of the fusion construct from mitochondria following mutation of the Fis1p TA leads to detection of sfGFP in the nucleus (C. Dunn, unpublished observations).

In the strain used for our selective scheme, a Ura+ phenotype requires greater Gal4-dependent transcriptional activation than is required for a His+ phenotype (Vidal *et al.* 1996b). Therefore, we reasoned that initial selection of TA mutants based on a His+ phenotype may provide informative mutations that weaken, but do not totally inhibit membrane association. We used mutagenic PCR to generate altered TAs within the context of a Gal4-sfGFP-Fis1 fusion protein. We then isolated four colonies that proliferated upon SMM-His medium containing 20mM 3-AT, yet exhibited diminished proliferation on SC-Ura medium when compared to cells expressing the Gal4-sfGFP-Fis1(ΔTA) polypeptide. Sanger sequencing of the region encoding the TA of Gal4-sfGFP-Fis1p within these colonies revealed one clone containing a V145E mutation (amino acid numbering provided in this study will correspond to that of the unmodified, full-length Fis1 protein; the necessary and sufficient region for mitochondrial association of Fis1p begins at amino acid L129), two clones containing a L139P mutation, and one clone harboring two mutations: L129P and V138A. Serial dilution assays (Figure 2A) confirmed that V145E and L139P provided a less than maximal, but still apparent Ura+ phenotype, with the V145E mutant allowing more rapid proliferation on medium lacking uracil than the L139P mutant. The L129P/V138A mutant provided a His+ phenotype, but could not drive uracil prototrophy, suggesting a less severe localization defect than that exhibited by the other two mutant TAs. Interestingly, the V145E mutation falls within the predicted MAD of the Fis1p TA (Figure S1A), consistent with poor accommodation of a negatively charged amino acid within this hydrophobic stretch of amino acids. Moreover, the Fis1p TA is predicted to be mostly alpha-helical in nature, and some evidence suggests that helicity is an important determinant of TA targeting to mitochondria (Wattenberg *et al.* 2007). Therefore, isolation of the potentially helix-disrupting L139P replacement by selection supports the need for TA helicity during mitochondrial targeting.

**Figure 2.**
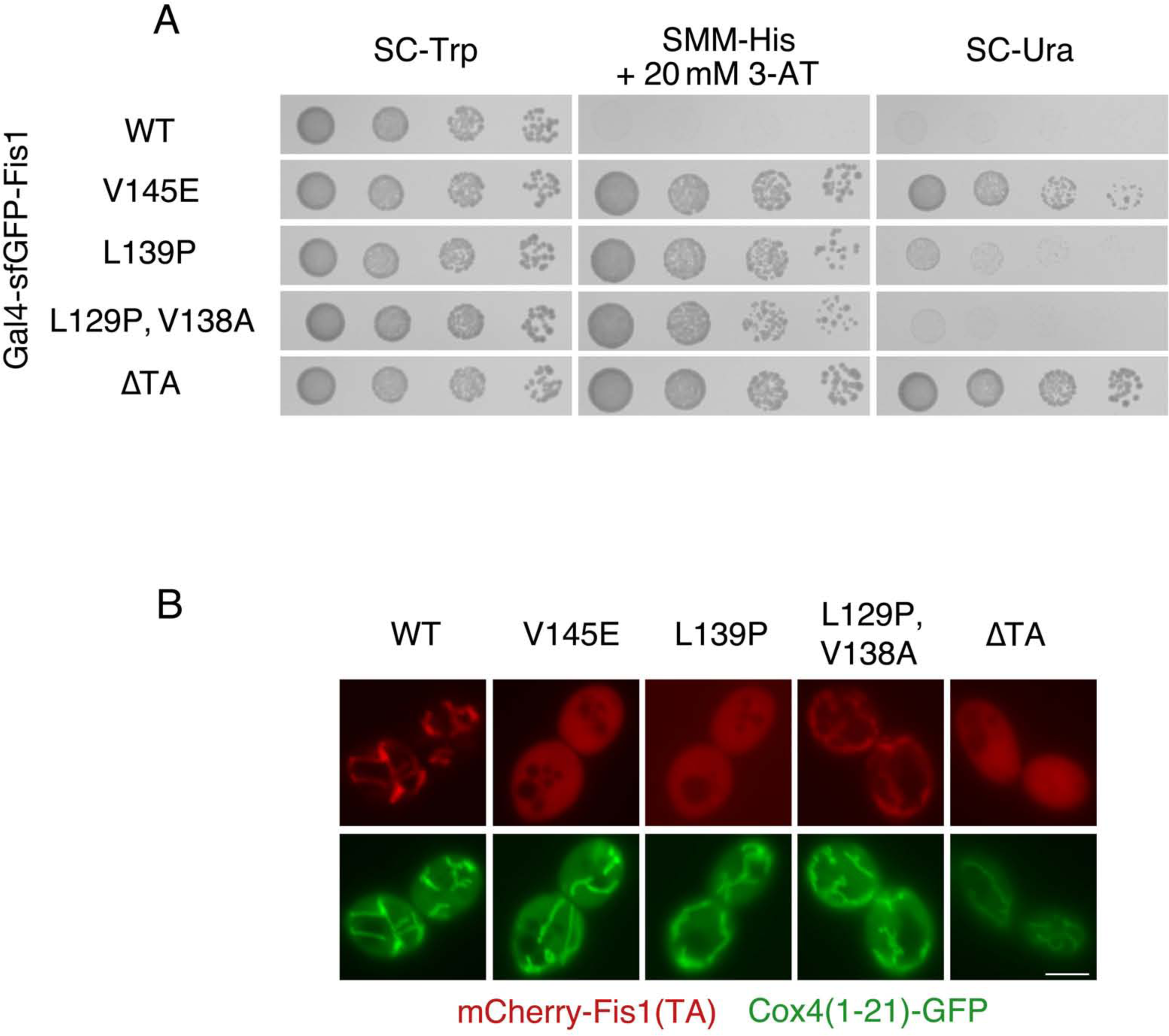
Selection for reporter activation by Gal4-sfGFP-Fis1p reveals TA mutations inhibiting mitochondrial localization. (A) Missense mutations within the Fis1p TA provide selectable marker activation. Strain MaV203 expressing Gal4-sfGFP-Fis1p from plasmid b100 or variants expressed from plasmids b128 (V145E), b129 (L139P), b130 (L129P, V138A), or b101 (ΔTA) were treated as in Figure 1C. pKS1 (vector) is also provided as a negative control. (B) Missense mutations within the Fis1p TA allow cytosolic and nuclear accumulation of a linked mCherry protein. mCherry fused to variants of the Fis1p TA were expressed in WT strain CDD961 from plasmids b109 (WT), b134 (V145E), b135 (L139P), b136 (L129P,V138A), or b252 (ΔTA) and visualized by fluorescence microscopy. Mitochondria were labelled with a mitochondria-targeted GFP expressed from plasmid pHS1. Scale bar, 5 μm.

To further examine whether isolated TA mutations affect mitochondrial OM targeting, we linked wild-type (WT) or mutant Fis1p TAs to the carboxyl-terminus of mCherry. To assess mitochondrial localization of these fusion proteins, mitochondria were specifically labelled with GFP targeted to mitochondria by the presequence of the Cox4 protein (Sesaki and Jensen 1999). Importantly, the TA of Fis1p lacks information required for mediating mitochondrial fragmentation (Habib *et al.* 2003), indicating that the localization of these constructs should not be influenced by interaction partners of full-length, inserted Fis1p.

The location of mCherry fused to Fis1p TAs was consistent with our genetic findings. V145E and L139P mutations in the Fis1p TA led to substantial cytosolic and nuclear accumulation of mCherry (Figure 2B). Moreover, the L129P/V138A TA, consistent with its weaker activation of Gal4 targets in our selection system, provided still discernable mitochondrial localization of the mCherry signal, but extraorganellar levels of this mutant fusion protein appeared to be increased compared to mCherry fused to the WT TA. These results suggest that our genetic approach is likely to allow recovery of mutations affecting the ability of the Fis1p TA to moor proteins to the mitochondrial outer membrane.

### A deep mutational scan for mutations potentially affecting Fis1p TA targeting

Buoyed by our initial isolation of TA mutations affecting Fis1p localization, we decided to take a more global approach to the analysis of the Fis1p TA. Using degenerate primers and recombination-based cloning in *S. cerevisiae*, we sought construction of a library consisting of all possible codons at every one of 27 amino acid positions within the TA of Gal4-sfGFP-Fis1p. We then allowed the pool of cells containing mutant TAs to divide four times under six possible culture conditions: no specific selection for *HIS3* or *URA3* reporter activation (SC-Trp, selecting only for plasmid maintenance), selection for *HIS3* activation at different 3-AT concentrations (SMM-Trp-His +0, 5, 10, or 20mM 3-AT), or selection for *URA3* activation (SC-Ura). Plasmid DNA was isolated from each pool of mutants, and TA coding sequences were amplified and subjected to next-generation sequencing. We focused our subsequent analysis only on clones within our pools harboring a WT TA sequence or encoding TAs with only a single amino acid change.

While all potential replacement mutations could not be detected within our starting library (Figure S2), and some biases did exist at each TA position, the vast majority of potential amino acid mutations were represented within our pool. 98.9% of potential amino acid replacements were identified in the starting pool cultured in SC-Trp, and 95.9% of TAs with single mutations were represented by at least 10 counts. Quantification of counts from all samples can be found in Table S1. When comparing the mutant pool cultured in SC-Trp with selection in SMM-Trp-His without added 3-AT, there was no appreciable difference in the relative abundance of most mutant TAs, including truncation mutations expected to totally prevent mitochondrial targeting of Gal4-sfGFP-Fis1p (Figure S3A). Such a result is consistent with ‘leaky’ expression of *HIS3* independent of Gal4-driven activation (Durfee *et al.* 1993). However, upon addition of 3-AT at concentrations of 5mM (Figure S3B), 10mM (Figure S3C), or 20mM (Figure 3) to medium lacking histidine, there were substantial shifts in the composition of the mutant pools toward specific amino acids, prompting further experiments that we describe below. The pool cultured in SC-Ura medium showed very strong selection for nonsense mutations within the TA (Figure S3D), but less prominent biases among amino acids. When considering our initial findings, in which recovery of uracil prototrophs by our genetic scheme led to a high recovery of frameshift and nonsense mutations, assessment of *HIS3* activation seems more informative regarding determinants of Fis1p TA targeting than competition assays performed in the more strongly selective medium lacking uracil. Independently of the primary amino acid sequence, the specific codons used to direct synthesis of a protein can affect that polypeptide’s translation rate, folding, or even synthesis of its transcript (Yu *et al.* 2015; Zhou *et al.* 2016). While the data are more ‘noisy’ due to a lack of representation of certain codons at each position, codons encoding the same amino acid generally acted in concert with one another within our selection scheme (Figure S4). Therefore, our focus remained upon the amino acid sequence of library variants rather than on codon sequence.

**Figure 3.**
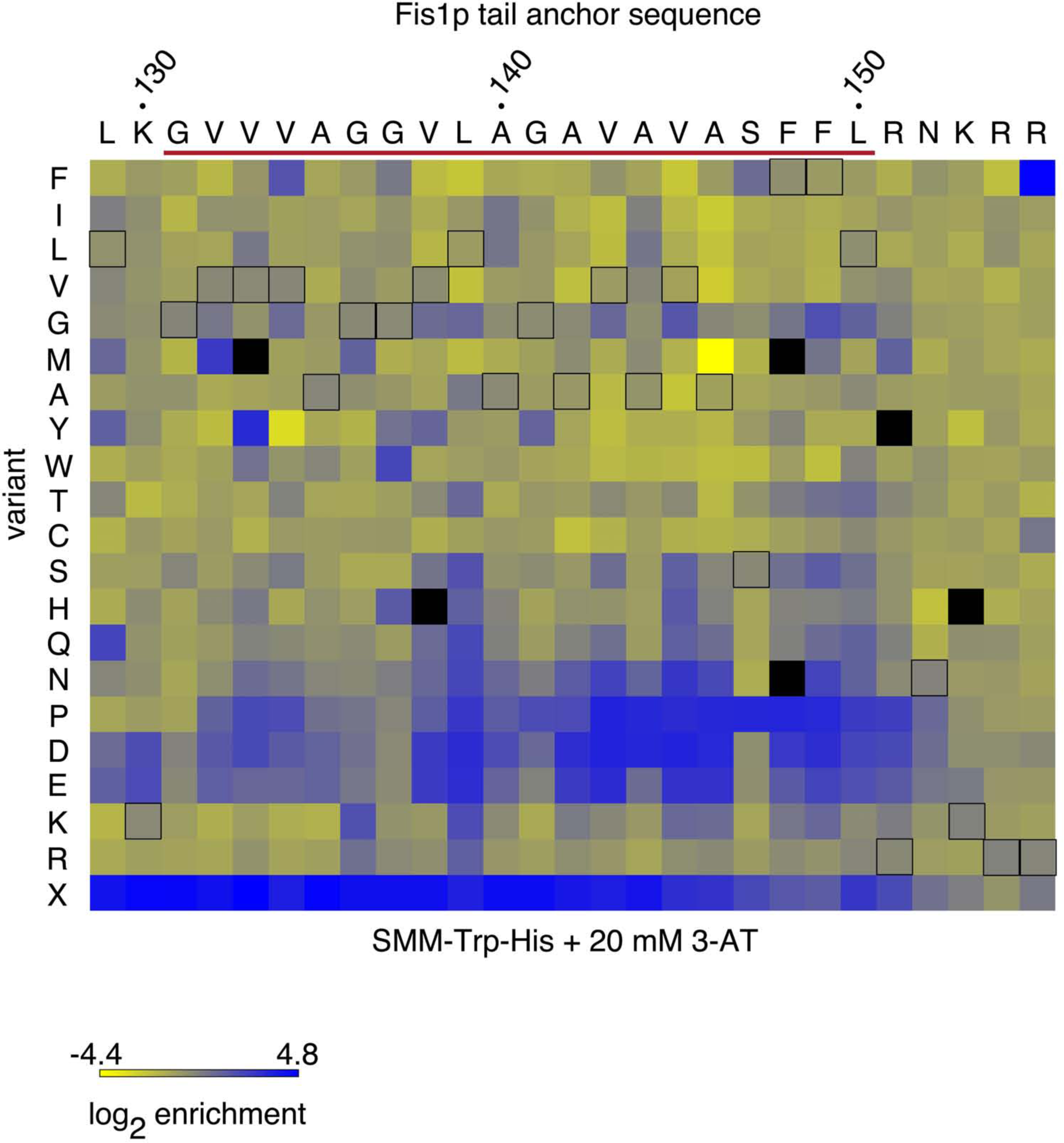
Global discovery of mutations within the TA of a Gal4-sfGFP-Fis1 fusion protein that allow Gal4-driven transcription. The log2 of enrichment values for each amino acid were calculated for each position following selection in SMM-Trp-His medium containing 20 mM 3-AT. Enrichment values are generated for individual amino acid positions within the TA, and not across positions. Black outlines denote the native amino acid for each position. Amino acid replacements not detectable under selective conditions are denoted by black, filled squares. The predicted MAD is indicated by a red line. ‘X’ represents substitution by a stop codon.

### Proline disruption of Fis1p tail anchor targeting

Previous analyses of various tail-anchored mitochondrial proteins similar in general structure to Fis1p suggested that no primary consensus sequence is required for TA insertion (Horie *et al.* 2002; Beilharz 2003; Rapaport 2003). While meaningful alignment of Fis1p TAs across species is difficult due to constraints in amino acid choice within hydrophobic domains and as a consequence of the apparently variable Fis1p TA length across species, only G131 (as pertains to the *S. cerevisiae* Fis1p sequence) might be considered highly conserved (Figure S1B). Our comprehensive analysis supports the idea that no consensus sequence within the Fis1p TA is necessary to achieve membrane insertion, since most amino acid replacements within the necessary and sufficient region required for Fis1p targeting, including at position G131, fail to lead to notable selectable reporter activation (Figure 3). Consequently, we focused our subsequent analysis on structural characteristics of the TA that might be most important for mitochondrial OM targeting.

The recovery of the L139P mutation during preliminary selection for Fis1p TA mutations indicated that proline may not be acceptable within the hydrophobic core of the Fis1p TA. Our deep mutational scan of the Fis1p TA in SMM-Trp-His + 20mM 3-AT (Figure 3) also strongly indicated that proline insertion across many positions disrupted mitochondrial TA localization. When focusing specifically upon those mutants that were in the top 75% most commonly counted variants in the starting pool (>126 counts) and enriched at least four-fold in SMM-Trp-His + 20mM 3-AT, 12 of 33 missense mutations within this set were proline replacements (Figure 4), further indicating failure of TA targeting following placement of proline at many TA positions.

**Figure 4.**
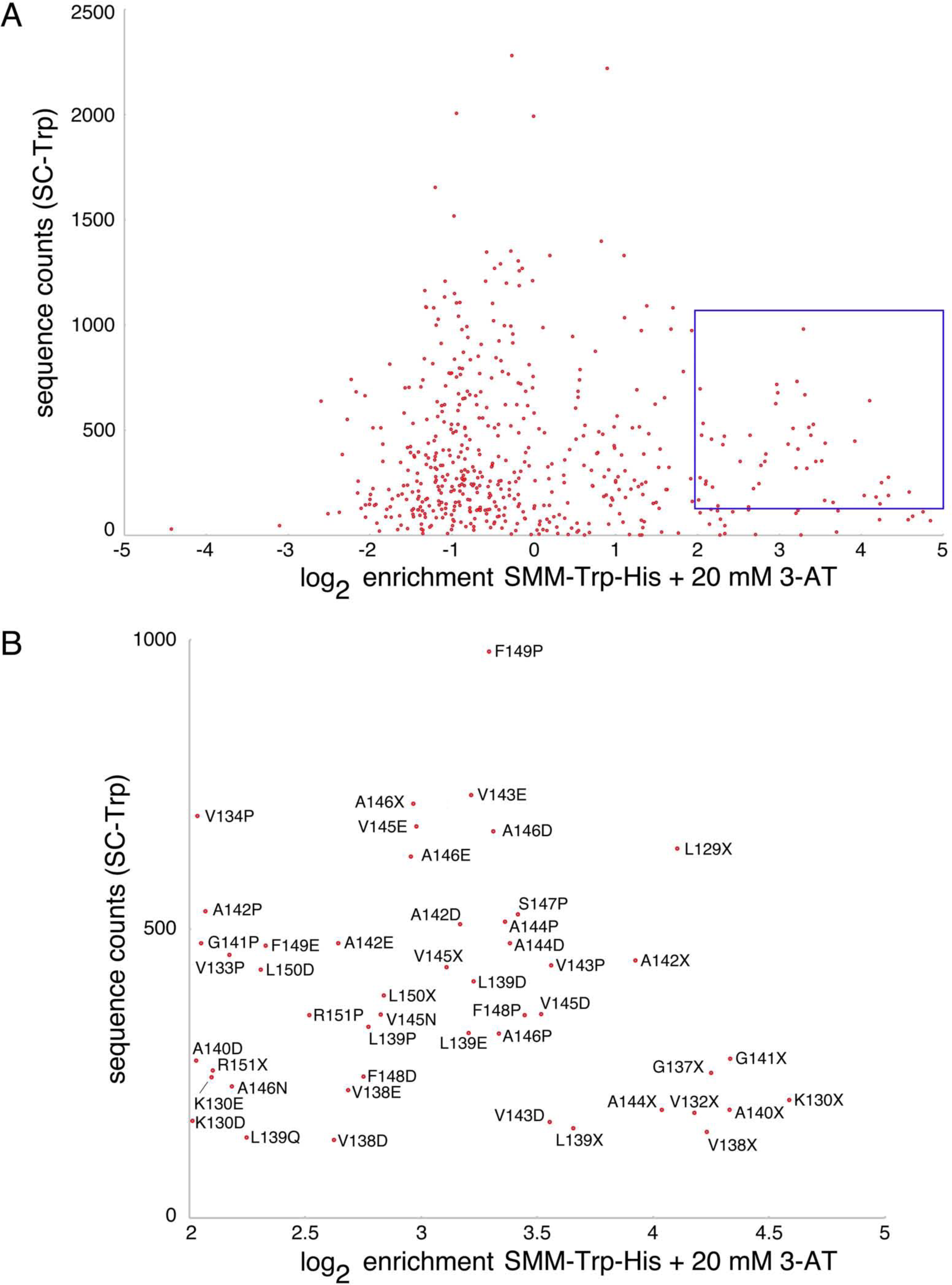
Identification of abundant Gal4-sfGFP-Fis1p clones which are highly enriched upon selection for Gal4-sfGFP-Fis1p nuclear translocation. (A) TA substitution mutations are plotted, with log2 enrichment values provided on the X-axis and sequence counts recovered from the starting pool (SC-Trp) provided on the Y-axis. Those replacement mutations that are within the top 75th percentile of mutant abundance in the starting pool and enriched at least four-fold following selection in SMM-Trp-His medium containing 20 mM 3-AT are highlighted in a blue box. (B) Expansion of the highlighted region in (A) showing specific TA mutations.

Subsequently, we carried out directed experiments to further examine poor accommodation of proline within the Fis1p TA. We further studied the L139P mutant that was initially isolated during selection for Fis1p TA targeting mutants, and we also generated four additional, individual proline replacements within Gal4-sfGFP-Fis1p and tested for Gal4-driven reporter activation. Newly constructed V134P, G137P, A140P, and A144P substitutions, consistent with our larger scale analysis (Figure 3 or Figure 4), provided ample proliferation on medium selective for *HIS3* activation (Figure 5A). Upon visualization of mCherry fused to these Fis1p TA mutants, V134P, L139P, A140P, and A144P replacements all clearly diminished mCherry localization to mitochondria (Figure 5B). Our results suggest that the secondary structure of the Fis1p TA is important for its function, and that disruption of helicity at many locations may make targeting to the mitochondrial OM unfavorable. We also noted, however, variability in the propensity of proline replacements to disrupt Fis1p TA targeting. Most prominently, the G137P substitution allowed apparently normal targeting to mitochondria, as assessed by Gal4p-driven reporter activation (Figures 3 and 5A) and by microscopic analysis (Figure 5B), potentially suggesting the existence of two separable, helical segments within the Fis1p TA rather than a single, monolithic helix.

**Figure 5.**
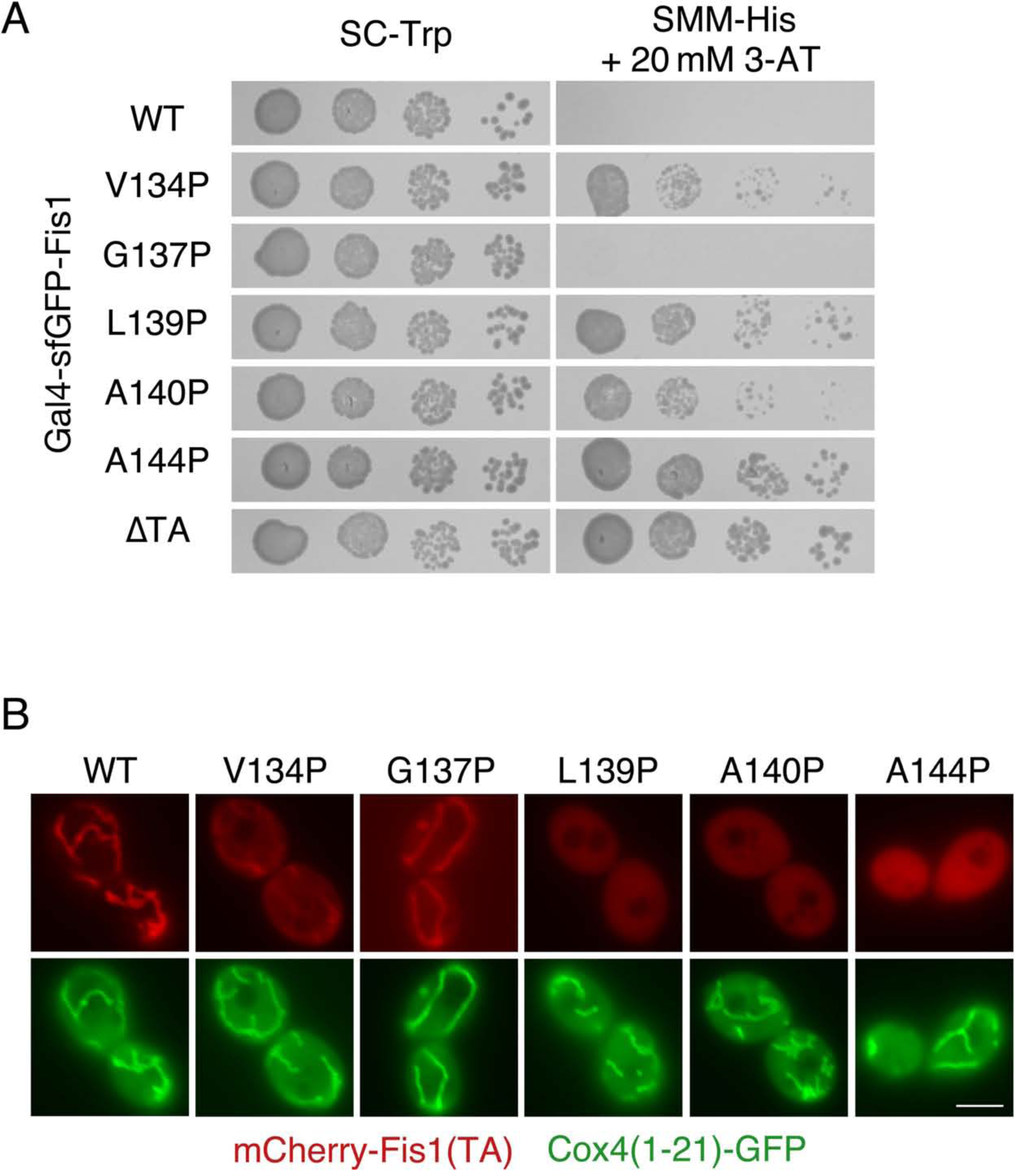
Proline substitution is acceptable at a discrete position within the Fis1p TA. (A) Replacement of specific amino acids within the TA of Gal4-sfGFP-Fis1p with proline can lead to Gal4-mediated selectable marker activation. Strain MaV203 expressing Gal4-sfGFP-Fis1p variants from plasmids b100 (WT), b188 (V134P), b189 (G137P), b129 (L139P), b190 (A140P), b296 (A144P), or b101 (ΔTA) was cultured in SC-Trp medium then spotted to SC-Trp or SMM-His + 20 mM 3-AT medium for 2 d. (B) TAs with specific proline replacements can reduce mitochondrial targeting of a linked fluorescent protein. Variants of the Fis1p TA fused to mCherry were expressed in WT strain CDD961 from plasmids b109 (WT), b208 (V134P), b209 (G137P), b135 (L139P), b210 (A140P), b211 (A144P) and examined, along with mitochondria-targeted GFP, as in Figure 2B. Scale bar, 5 μm.

We considered the possibility that targeting of these mutant TAs is not impeded, but rather a lack of stability within the lipid bilayer leads to ejection from the mitochondrial OM by a quality control mechanism. Recently, the yeast Msp1 protein and its human ortholog ATAD1, have been identified as potential ‘extractases’ that can remove improperly folded or mislocalized proteins from the mitochondrial OM (Okreglak and Walter 2014; Chen *et al.* 2014). We tested whether any of tail-anchored fluorescent proteins containing proline replacements and not strongly localized to mitochondria could recover mitochondrial localization in cells deleted of Msp1p. However, deletion of Msp1p did not lead to relocalization of any tested mutant mCherry-TA fusion protein to mitochondria (Figure S5A), providing no evidence for proper targeting, then subsequent removal, of assayed Fis1p TAs harboring proline replacements.

In mammalian cells, the ubiquilin protein UBQLN1 can bind to tail-anchored proteins that fail to target mitochondria, expediting their degradation (Itakura *et al.* 2016). We wondered whether the yeast UBQLN1 orthlog Dsk2p (Chuang *et al.* 2016) might bind to mutant Fis1p TAs, sequestering them in the cytosol and preventing mitochondrial insertion. However, deletion of Dsk2p did not change the cellular location of mislocalized mCherry-TA fusion proteins carrying proline substitutions (Figure S5B).

### The positively charged carboxyl-terminus of the tail anchor allows both efficient and specific targeting to mitochondria

Analysis of the data from our deep mutational scan suggested that nonsense mutations throughout much of the TA can allow Gal4-sfGFP-Fis1p to move to the nucleus and activate transcription (Figures 3, 4, and S3). Stop codons placed within the highly charged RNKRR pentapeptide near the carboxyl-terminus of Fis1p, however, seem to permit some membrane localization, as reported by proliferation rates in selective medium. Therefore, we examined the behavior of a R151X mutant, which lacks all charged amino acids following the predicted MAD (X represents mutation to a stop codon). Supporting partial localization to a cellular membrane, the R151X mutant of Gal4-sfGFP-Fis1p did not activate Gal4-controlled expression of *HIS3* to the same extent of a Gal4-sfGFP-Fis1p construct lacking the entire TA (Figure 6A). Consistent with those results, the R151X TA directed mCherry to intracellular organelles (Figure 6B). However, along with some apparent mitochondrial association, the R151X TA was also clearly localized to the ER (Figure 6C). Interestingly, ER localization of mCherry fused to the R151X TA was not completely dependent upon Get3p, a receptor for ER tail-anchored proteins (Schuldiner *et al.* 2008), suggesting the existence of an alternative pathway for localization of the R151X TA to the ER (Figure S6A and S6B). Similarly, R151X TA could be targeted to the ER in *get2Δ* mutants, lacking a receptor for several ER-directed tail-anchored proteins (Figure S6C). However, puncta that might indicate clustering with Get3p (Schuldiner *et al.* 2008) were apparent in some *get2Δ* cells, raising the unexplored possibility that Get3p can bind at least some portion of the cytosolic pool of R151X TA.

**Figure 6.**
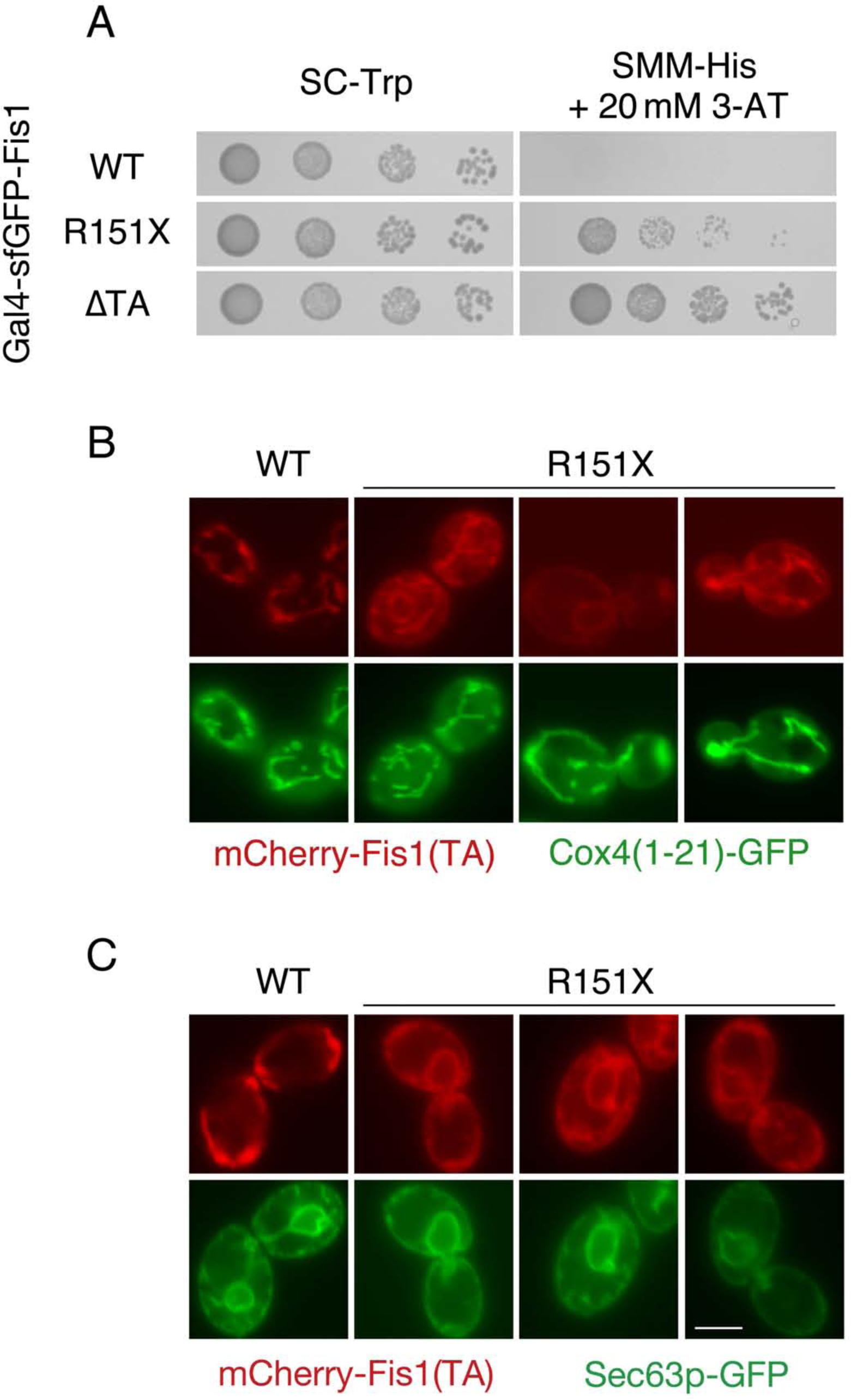
The positively charged carboxyl-terminus of the Fis1p TA is important for specific localization to and insertion at the mitochondrial outer membrane. (A) Deletion of the final five amino acids from the Fis1p TA permits transcriptional activation by Gal4-sfGFP-Fis1p. Strain MaV203 harboring plasmids b100 (WT), b253 (R151X), or b101 (ΔTA) was treated as in Figure 5A. (B) Removal of the last five amino acids from the Fis1p TA allows mislocalization to the secretory system. Strain CDD961 expressing mCherry fused to the WT Fis1p TA from plasmid b109 or expressing mCherry linked to a truncated Fis1p TA (R151X) from plasmid b254 was evaluated as in Figure 2B. (C) Strain CDD961 was cured of plasmid pHS1. The resulting strain was transformed with plasmid pJK59 to label ER by expression of Sec63p-GFP, then transformed with either plasmid b109 or plasmid b254 to localize the WT and R151X TAs and examined by fluorescence microscopy. Scale bar, 5 μm.

Together, our results demonstrate that the charged amino acids at the carboxyl-terminus of the TA provide organelle specificity, yet are not totally required for membrane localization. These findings are consistent with previous results reporting that positively charged amino acids following the MAD of Fis1p allow targeting of TAs specifically to mitochondria rather than ER (Isenmann *et al.* 1998; Kuroda *et al.* 1998; Borgese *et al.* 2001; Stojanovski *et al.* 2004), perhaps by decreasing total TA hydrophobicity, a factor important for determining the final location of a TA (Beilharz 2003; Wattenberg *et al.* 2007).

Since the R151X variant of Gal4-sfGFP-Fis1p activated Gal4-driven reporters, yet was at least partially localized to the ER, we wondered if ER localization of any protein fused to Gal4 might similarly lead to Gal4-driven transcription due to the physical continuity between the ER and nuclear envelope. Therefore, we examined the TA of the human FIS1 protein (hFIS1), since full-length hFIS1 can localize to ER in *S. cerevisiae* (Stojanovski *et al.* 2004). Indeed, we found that mCherry fused to the hFIS1 TA was poorly localized to mitochondria (Figure S7A) and abundant at the ER (Figure S7B). However, a fusion protein consisting of Gal4 fused to the TA of hFIS1 did not provide *HIS3* activation (Figure S7C), indicating that the hFIS1 TA is quantitatively membrane-targeted in *S. cerevisiae* and that activation of Gal4-dependent reporters upon removal of the positively charged carboxyl-terminus from Gal4-sfGFP-Fis1p is unlikely to be a consequence of ER localization.

Due to the mislocalization of the hFIS1 TA, we then investigated the possibility that other mitochondrial TA proteins from human would be targeted improperly in *S. cerevisiae*. We fused mCherry to the TA of human BAX, a region that is sufficient for insertion at the mammalian mitochondrial OM (Schinzel *et al.* 2004). While mCherry signal was diminished in comparison with other mCherry fusion proteins examined in this study and expressed under the same promoter, mCherry fused to the BAX TA was properly targeted to mitochondria (Figure S7D). Gal4 fused to the BAX TA did not activate selectable reporters (C. Dunn, unpublished results), further indicating effective mitochondrial targeting mediated by the BAX TA.

### Extension or reduction of Fis1p TA length does not affect targeting to mitochondria

Targeting of tail-anchored proteins to specific membranes has been suggested to depend, at least in part, upon the specific length of the MAD within the TA (Isenmann *et al.* 1998; Horie *et al.* 2002). We reasoned that the region within the MAD at which prolines do not strongly disrupt mitochondrial targeting, as defined by our analysis, may be amenable to the insertion or deletion of new amino acids, thereby allowing us to test the relationship between Fis1p TA length and mitochondrial targeting. We inserted one (∇1A), two (∇2A), or three (∇3A) additional alanines between A135 and G136 within the TA of Gal4-sfGFP-Fis1p, but none of these mutant constructs led to apparent *HIS3* (Figure 7A) activation. We then analyzed the location of mCherry fused to a Fis1p TA carrying these same insertions. All constructs were localized properly to mitochondria (Figure 7B).

**Figure 7.**
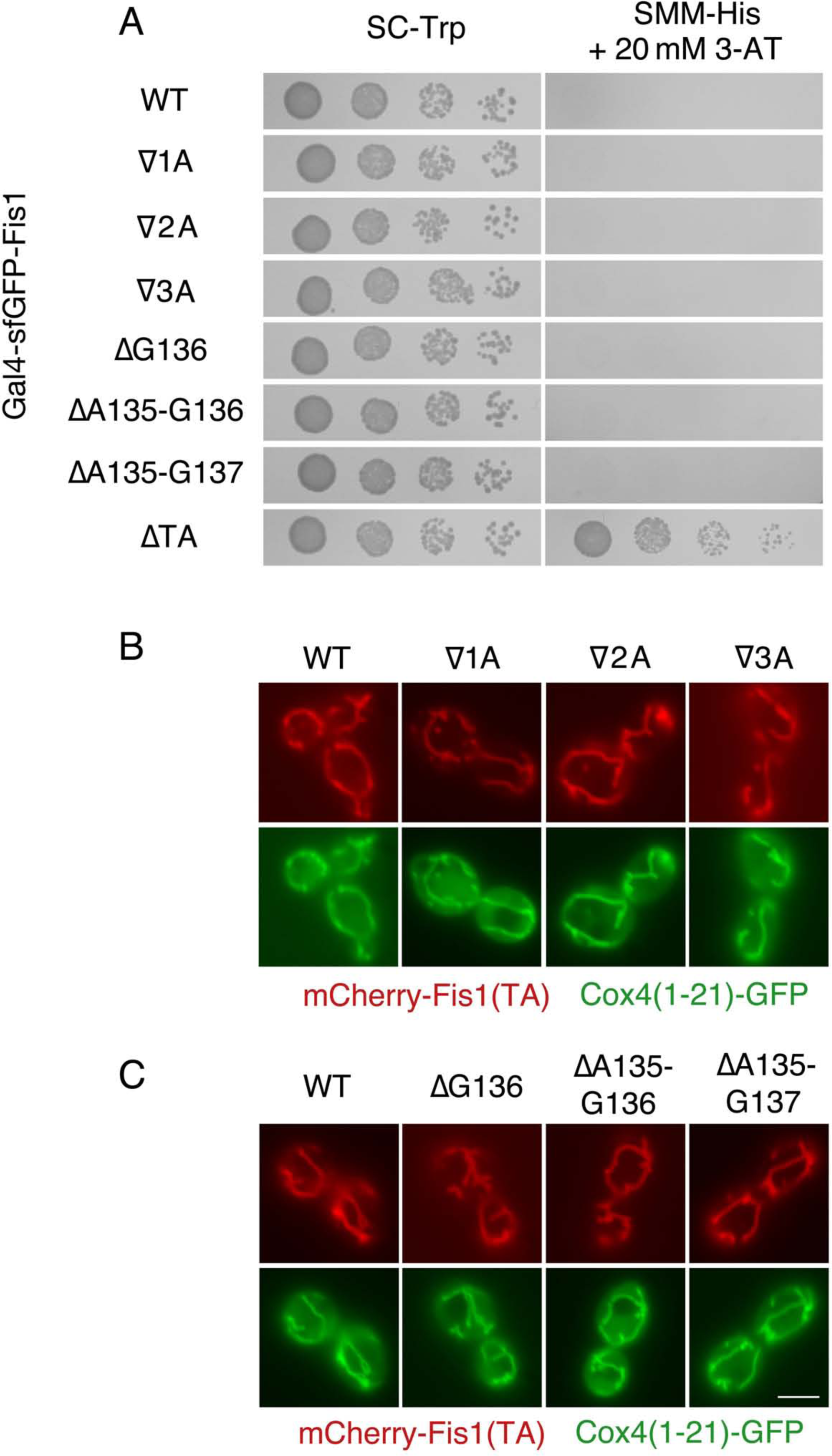
Targeting of the Fis1p TA is not dependent upon a specific TA length. (A) Deletion of up to three amino acids or insertion of up to three amino acids does not allow Gal4-sfGFP-Fis1p to activate transcription. MaV203 cells expressing Gal4-sfGFP-Fis1p variants from plasmids b229 (∇1A), b230 (∇2A), b231 (∇3A), b226 (ΔG136), b227 (ΔA135-G136), b228 (ΔA135-G137), or b101 (ΔTA) were treated as in Figure 5A. (B) mCherry fused to a Fis1p TA containing an insertion of up to three amino acids in length localizes properly to mitochondria. Strain CDD961 expressing mCherry-TA fusions from plasmids b109 (WT), b235 (∇1A), b236 (∇2A), or b237 (∇3A) was visualized as in Figure 2B. (C) mCherry fused to a Fis1p TA deleted of up to three amino acids is properly targeted to mitochondria. Strain CDD961 expressing mCherry-TA fusions from plasmids b109 (WT), b232 (ΔG136), b233 (ΔA135-G136), or b234 (ΔA135-G137) was examined as in Figure 2B. Scale bar, 5 μm.

Next, we deleted one (ΔG136), two (ΔA135-G136), or three (ΔA135-G137) amino acids within the Fis1p MAD and performed similar assays. Like our insertion mutants, deletion mutants were apparently targeted to a membrane, as assessed by Gal4-driven reporter transcription (Figure 7A). Moreover, mCherry remained localized to mitochondria when up to three amino acids were deleted (Figure 7C).

Disruption of the ER-localized Spf1 protein reduces the contrast in ergosterol content between the ER and the mitochondrial OM (Krumpe *et al.* 2012). Consequently, TAs normally localized to mitochondria are mistargeted to the ER upon Spf1p deletion. The sterol concentration of membranes can determine bilayer thickness (Dufourc 2008), raising the possibility that insertions or deletions may allow mitochondrial TAs to once again prefer mitochondrial OM targeting over ER localization in a background lacking Spf1p. However, mCherry linked to insertion or deletion mutants of the Fis1p TA remained poorly targeted to mitochondria (Figure S8A) and prominently localized to the ER (Figure S8B) in mutants lacking Spf1p.

Together, our results demonstrate that the Fis1p TA is properly localized to mitochondria even when its length is substantially altered.

### Positively charged amino acids are more acceptable than negatively charged amino acids within the predicted transmembrane domain of the Fis1p tail anchor

Our deep mutational scan of the Fis1p TA demonstrated that Gal4-sfGFP-Fis1p was generally able to activate gene expression when aspartate or glutamate was placed within the MAD (Figure 3). In fact, upon examination of those amino acid replacements found within the top three quartiles of counts in the initial library and also enriched at least four-fold upon culture in SMM-Trp-His + 20mM 3-AT, 18 of 33 missense mutations were aspartate or glutamate substitutions (Figure 4). We were surprised to find that placement of positively charged arginine or lysine residues appeared to be much more acceptable within the MAD of Fis1p than aspartate or glutamate; none of the amino acid substitutions within this high-count, high-enrichment set were by lysine or arginine.

To further pursue the possibility that positively charged amino acids can be accommodated within the Fis1p MAD, we mutated four amino acids within the hydrophobic stretch of the Fis1p TA to aspartate, glutamate, lysine, or arginine. Specifically, we generated amino acid replacements at positions V132, A140, A144, or F148, then retested these mutants under selection for Gal4-sfGFP-Fis1p transcriptional activity. The results from our global analysis were verified, with aspartate and glutamate mutations providing stronger reporter activation than lysine and arginine mutations (Figure 8A). Only the A144D mutation provided sufficient Gal4 activation for proliferation on medium lacking uracil (Figure S9A) after two days of incubation, suggesting a very severe TA localization defect caused by this TA mutation. We noted that these mutant Gal4-sfGFP-Fis1 constructs exhibit altered behavior at different temperatures. For example, lysine and arginine substitutions at positions A140, A144, or F148 led to reduced proliferation at 37°C under conditions selective for *HIS3* activation (Figure S9B) when compared to the same TA substitutions assessed at 18°C (Figure S9C) or 30°C (extended incubation, Figure S9D). This outcome is consistent with the idea that altered phospholipid dynamics at different temperatures may lead to consequent changes in TA insertion efficiency (de Mendoza and Cronan 1983).

**Figure 8.**
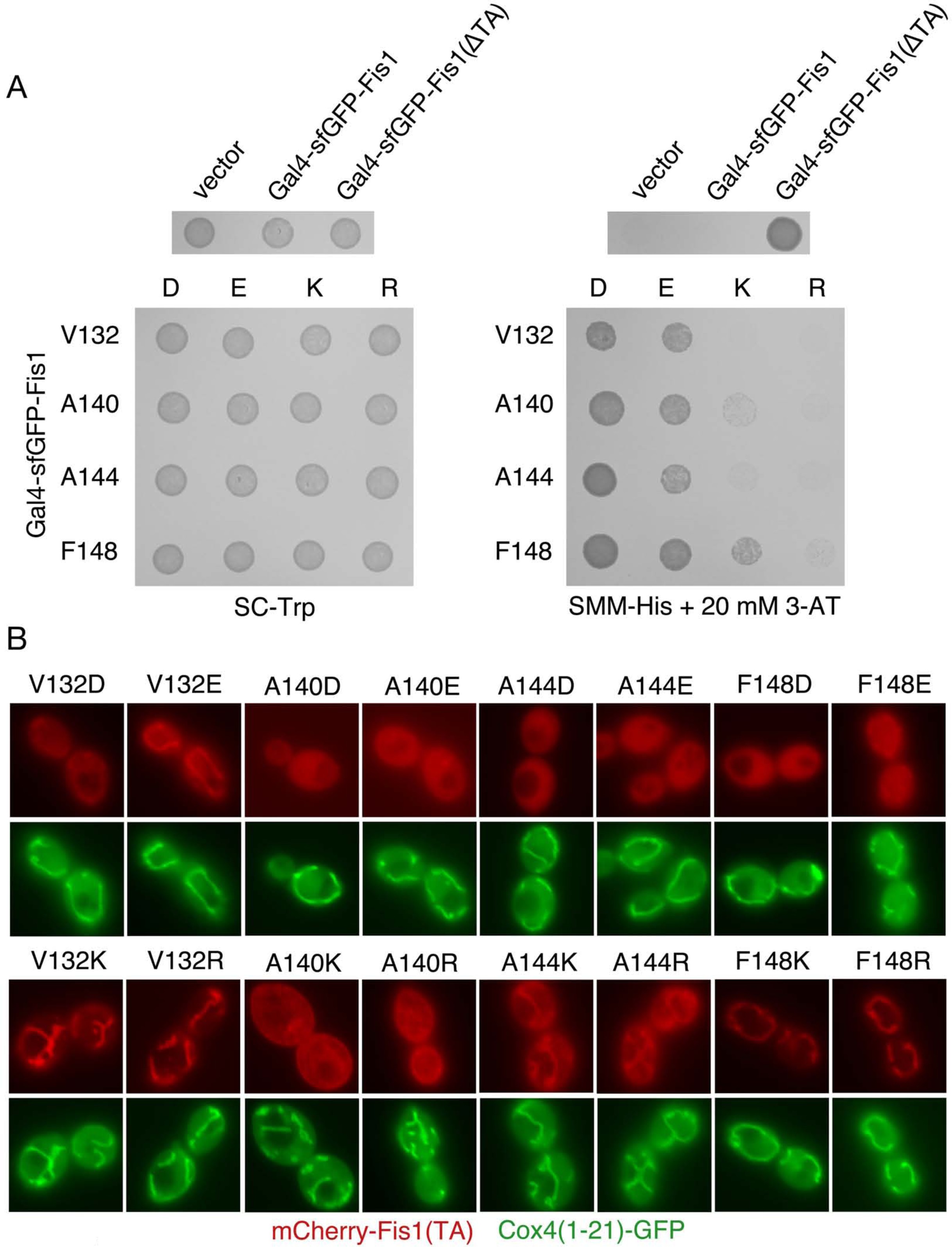
Fis1p TA targeting is hindered to a greater extent by inclusion of negatively charged amino acids in the MAD than by positively charged amino acids. (A) Negative charges allow higher transcriptional activity than positive charges when placed at specific positions within the Gal4-sfGFP-Fis1p TA. Strain MaV203 was transformed with plasmids pKS1 (vector), b100 (Gal4-sfGFP-Fis1), b101 [Gal4-sfGFP-Fis1(ΔTA)], or plasmids encoding the indicated charge replacements within the Fis1p TA (plasmids b173-b187 and b295). The resulting transformants were spotted to SC-Trp medium (1 d, 30°C) or SMM-His + 20mM 3-AT medium (2 d, 30°C). (B) mCherry-TA localization is disrupted more severely by negatively charged amino acids within the MAD than by positively charged amino acids. Strain CDD961 was transformed with plasmids (b192-b207) expressing mCherry linked to Fis1p TAs harboring the indicated substitutions. Cells were visualized as in Figure 2B. Scale bar, 5 μm.

We then tested the ability of these Fis1p TAs containing charge substitutions to promote mitochondrial localization of mCherry. At V132 and F148, positions within the MAD nearer to the putative water-lipid bilayer interface, mutation to positively charged amino acids allowed abundant localization to mitochondria (Figure 8B). In contrast, mutation to negatively charged amino acids clearly hindered mitochondrial targeting. We noted that F148D and F148E replacements hampered mitochondrial localization more severely than V132D and V132E replacements, consistent with phenotypic testing of Gal4-sfGFP-Fis1 fusion proteins. At position A144, lying more deeply within the predicted MAD, all charge mutations inhibited mCherry targeting, yet A144K and A144R substitutions allowed some mCherry localization at mitochondria while A144D and A144E substitutions appeared undetectable at mitochondria. Finally, no mitochondrial signal was apparent for any of the charge mutants tested at position A140 of the Fis1p TA. However, A140K and A140R mutants differed from A140D and A140E mutants by localizing to other membranes within the cell, including the plasma membrane, rather than providing a diffuse cytosolic signal. Msp1p removal did not permit relocalization of tail-anchored fluorescent proteins to mitochondria (Figure S5C), supporting the idea that charge replacements within the Fis1p TA lead to a failure of association with the OM rather than enhanced removal from mitochondria. Removal of UBQLN1 ortholog Dsk2p also had no discernable effect on mCherry-TA mutant localization (Figure S5D).

Negligible Fis1p activity at mitochondria is apparently sufficient to promote mitochondrial fission (Habib *et al.* 2003; Krumpe *et al.* 2012), suggesting that even minimal localization and related functionality at the OM would be detectable by functional assays. Interestingly, Fis1p variants harboring charged residues, positive or negative, at positions V132, A144, or V148, with the exception of Fis1p carrying the A144D mutation, provided at least some Fis1p function, as indicated by microscopic (Figures S10A and S10B) and genetic (Figure S10C) assays. Such a result is consistent with the finding that only the A144D charge mutant provides sufficient *URA3* activation for rapid proliferation on medium lacking uracil (Figure S9A).

Together, our results demonstrate that positively charged amino acids within the MAD can better promote Fis1p localization than negatively charged amino acids, but that even negatively charged amino acids can be accommodated within the MAD and lead to a detectable level of mitochondrial targeting.

## DISCUSSION

Using a deep mutational scanning approach, we explored structural characteristics of the Fis1p TA important for targeting to the mitochondrial outer membrane. To our knowledge, this work is the first application of this technique to the study of a eukaryotic organelle targeting signal. Deep mutational scanning, when coupled to an effective method of screening or selection, is very cost‐ and time-effective (Araya and Fowler 2011; Fowler and Fields 2014; Boucher *et al.* 2014). Mutant library generation, subsequent pool selection, and next-generation sequencing can be completed in just a few months. Consequently, this approach generates far more useful data over a shorter duration than, for example, alanine scanning mutagenesis or low-throughput genetic selection followed by Sanger sequencing. Deep mutational scanning has recently been applied successfully to other areas of study, such as membrane protein insertion within bacteria (Elazar *et al.* 2016), tumor suppressor structure and function (Starita *et al.* 2015), and the relationship between a polypeptide’s evolutionary path and its present fitness (Hietpas *et al.* 2011; Melamed *et al.* 2013). As is true for most genetic screens and selections, further experimentation, as performed here by testing mutant Fis1p TAs in microscopic and functional assays, is required in order to make full use of a larger-scale dataset. We also note that some mutations affecting specific insertion of the Fis1p TA at mitochondria may have been missed during our global analysis: transcription driven by Gal4-sfGFP-Fis1p indirectly reports on TA targeting and may be affected by mutant stability, potential sequestration at other intracellular membranes, or ability of the fusion protein to enter and function within the nucleus.

### In vivo evidence of the ability of positively charged amino acids to “snorkel” their side chains to surface of the lipid bilayer

Because there is a high energetic barrier to placing any charged residue into a lipid bilayer (Cymer *et al.* 2015), we were initially surprised to find that positively charged amino acids within the MAD of the Fis1p TA promoted mitochondrial targeting far better than negatively charged amino acids. However, it has been suggested that lysine and arginine within a MAD can “snorkel,” or integrate the uncharged portion of their side chain into the hydrophobic milieu of the lipid bilayer while locating positive charges near polar head groups at the interface between the membrane and aqueous environment (Gavel *et al.* 1988; Segrest *et al.* 1990). Some phospholipid head groups carry a net negative charge, potentially providing further favorability to snorkeling lysine and arginine. The shorter hydrophobic portion of aspartate and glutamate, however, may not permit the side chain to easily reach the membrane interface to remove the negative charge from the hydrophobic environment, and if they do reach the lipid bilayer surface, charge repulsion might make such a conformation unfavorable.

Most experimental support for amino acid snorkeling has been obtained using *in vitro* approaches or has been gathered during structural studies (Monné *et al.* 1998; Long *et al.* 2005; Kim *et al.* 2012; Öjemalm *et al.* 2016), and little *in vivo* evidence has been reported in support of this phenomenon. Our comprehensive study of the Fis1p TA, a region dedicated only to the process of membrane integration (Habib *et al.* 2003; Kemper *et al.* 2008), strongly supports the ability of lysine or arginine to be accommodated by snorkeling at numerous positions within the Fis1p MAD. We note that snorkeling, if operative for positive charges within the Fis1p TA, may not be permitted within the context of all mitochondrial TAs: replacement of S184 within the BAX TA by lysine does not seem to permit mitochondrial localization of this protein (Nechushtan *et al.* 1999). We also note that while the Fis1p TA is typically modelled as bitopic, or reaching through the mitochondrial outer membrane from the cytosol to the intermembrane space, the possibility that the Fis1p TA lies side-long in the outer membrane has not been ruled out. However, snorkeling is thought to be possible for a MAD found in either the monotopic or the bitopic configuration (Strandberg and Killian 2003).

Further comprehensive mutational scans of MADs may further substantiate the concept of snorkeling. Interestingly, since those Fis1p TAs mutated to contain positively charged amino acids within the MAD were often not targeted to the OM with full efficiency, one might imagine a scenario in which dual localization of a protein with a single MAD to cytosol and to intracellular membranes can be easily evolved by the appearance of positively charged amino acids at positions previously lacking such charges. Moreover, some prediction methods for MADs might be considered overly conservative, especially for prediction programs emphasizing amino acid charge. Therefore, the accumulating evidence of amino acid snorkeling should prompt the development of improved algorithms that explicitly consider positively charged amino acids to be acceptable within certain positions of a predicted MAD.

### The membrane-associated domain of the Fis1p tail-anchor may consist of two separable segments

Computational analyses suggest that the Fis1p TA is mostly alpha-helical in nature (Buchan *et al.* 2013; Drozdetskiy *et al.* 2015). We found that proper localization of the Fis1p TA likely requires its predicted alpha-helicity within the MAD, since proline, which is known to break or kink helices (Senes *et al.* 2004), profoundly disrupts targeting when substituted at many positions throughout this hydrophobic region. However, we found that replacement by proline is more acceptable at a specific location, G137, than proline mutations found at many other locations within this region, potentially indicating that the Fis1p MAD is bipartite in nature. Further supporting a bipartite structure of the Fis1p MAD, insertion of new amino acids between A135 and G136 did not apparently affect mitochondrial TA targeting. Moreover, mutations toward the carboxyl-terminal end of the Fis1p MAD appear to affect mitochondrial targeting more drastically, as reported by deep mutational scanning, than mutations toward the amino terminus of the transmembrane segment. Previous analysis of the rat OMP25 TA also support a bipartite structure of the MAD, with higher sensitivity to mutation nearer to the carboxyl-terminal end of this hydrophobic stretch (Horie *et al.* 2002). In addition, prolines are found within the MAD of the mammalian OMb and OMP25 TAs. These results suggest that those prolines might demarcate the boundary between distinct structural regions of the targeting sequence. On the other hand, prolines within a single helical segment may simply be more easily housed within an alpha-helix when buried deep in the lipid bilayer (Li *et al.* 1996; Senes *et al.* 2004) and may not reflect two separable MAD segments. If this is the case, prolines found in mitochondrial TAs might indicate the portion of the TA found at the midpoint of the OM.

Glycine is not preferred within alpha-helices (Chou and Fasman 1974; O’Neil and DeGrado 1990) as a consequence of its conformational flexibility. However, our deep mutational scan does not indicate reduced membrane targeting when most amino acids within the Fis1p TA are individually mutated to glycine. This might be surprising in light of the pronounced effects provided by several proline replacement mutations throughout this domain. Yet, glycine may not be as disruptive for alpha-helices found within a lipid bilayer environment when compared with alpha-helices of soluble proteins, due to better intra-helical hydrogen bonding within the hydrophobic environment of the membrane (Dong *et al.* 2012). Indeed, four glycines already exist within the *S. cerevisiae* Fis1p TA, and the TAs of Fis1p orthologs are also littered with glycines (Stojanovski *et al.* 2004) and (Figure S1B), further indicating that glycines are less disruptive of the Fis1p TA than prolines. Interestingly, GXXXG motifs, and other similarly spaced small amino acids like alanine and serine, can promote helix packing within lipid bilayers (Russ and Engelman 2000; Senes *et al.* 2004; Gimpelev *et al.* 2004). However, our findings indicate that the sole Fis1p GXXXG motif and a nearby AXXXA motif do not play a significant role in targeting of the Fis1p TA to membrane. Furthermore, a GXXXXG motif may mediate multimerization of the mitochondrial OM protein Bcl-XL (Ospina *et al.* 2011), and such a motif can be found within the *S. cerevisiae* Fis1p and several of its orthologs. However, the GXXXXG motif also seems to have little to no role in insertion, at least as determined by the ability of Gal4-sfGFP-Fis1p to activate gene expression.

### Does a machinery dedicated to mitochondrial tail anchor insertion exist?

So far, no cellular machinery dedicated to the insertion of mitochondrial tail anchored proteins has been revealed. Evidence supporting the existence of such a machinery includes saturable import of mitochondrial TA-containing proteins in mammalian cells (Setoguchi *et al.* 2006), potentially indicating a finite number of binding sites for mitochondrial TAs. Furthermore, the TOM complex has been reported to assist in insertion of full-length BAX into mitochondria (Ott *et al.* 2007; Cartron *et al.* 2008; Colin *et al.* 2009). Consistent with the need for a mitochondrial TA translocation machinery, the hFIS1 TA localizes specifically to mitochondria in human cells (Suzuki *et al.* 2003), but cannot effectively localize mCherry to *S. cerevisiae* mitochondria, possibly suggesting evolutionary divergence and a structural mismatch between the hFIS1 TA and any putative yeast TA translocation apparatus. However, we note that not all human mitochondrial TAs fail to be imported at the proper organelle in yeast, since our genetic and microscopic results indicate that the human BAX TA can be targeted to yeast mitochondria.

Other, perhaps more abundant evidence supports the idea that mitochondrial tail-anchored proteins similar in structure to Fis1p do not require a translocation machinery and can spontaneously insert into the OM. First, the MAD of the TA is protected from chemical modification upon exposure to lipid vesicles devoid of protein (Kemper *et al.* 2008), suggesting that the Fis1p TA can insert into lipid bilayers without assistance. Moreover, blockade or destruction of the general insertion pore for mitochondrial proteins, associated receptors, or other outer membrane protein biogenesis machinery such as the SAM complex or the MIM complex did not prevent Fis1p insertion at yeast mitochondria (Stojanovski *et al.* 2007; Kemper *et al.* 2008; Sinzel *et al.* 2016). Tail-anchored proteins also appear to have the ability to spontaneously and rapidly insert into mammalian mitochondria without the need for the TOM complex or soluble cytosolic chaperones (Setoguchi *et al.* 2006), although cytosolic chaperones potentially play a role in maintaining solubility of tail-anchored proteins while they are *en route* to mitochondria (Itakura *et al.* 2016). Further supporting the absence of a translocation machinery dedicated to TA insertion, a large-scale screen for proteins required for proper localization of mitochondrial tail-anchored proteins uncovered no putative translocon components (Krumpe *et al.* 2012). We note that Fis1p is also localized to peroxisomes (Kuravi *et al.* 2006), suggesting that any machinery that allows Fis1p TA insertion would potentially be shared by both mitochondria and peroxisomes. However, no dual-localized translocation machinery has yet been identified.

While a fraction of Fis1p can be targeted to the peroxisome, we found that detection of mutant Fis1p TAs in the cytosol and nucleus by microscopic and genetic assays is not driven by a specific failure of peroxisomal insertion. mCherry fused to the wild-type Fis1p TA is not evident in the cytosol or nucleus upon severe disruption of peroxisomal biogenesis (Hettema *et al.* 2000) by deletion of the peroxisomal membrane protein import components Pex3p (Figure S11A) or Pex19p (C. Dunn, unpublished observations). Moreover, reporter activation by Gal4-sfGFP-Fis1p containing an unmutated TA is not increased when *PEX3* is removed (Figure S11B), and the relative level of Gal4p-driven reporter activation among proline substitution mutants (Figure S11C) or charge substitution mutants (Figure S11D) did not differ between *PEX3* and *pex3Δ* cells.

If a TA insertion machinery does exist at the mitochondrial OM, loss-of-function mutations affecting this machinery would presumably be recovered by the application of our genetic selection scheme. Moreover, we expect that refined analysis of other organelle targeting signals and membrane insertion sequences will be accomplished by applying the deep mutational scanning approach outlined in this study.

## MATERIALS AND METHODS

### Yeast strains and plasmids

Details of strains used in this study are provided in Table S2. Plasmid acquisition details and associated references, as well as details of plasmid construction, are found in Table S3. Oligonucleotides used in this study are listed in Table S4.

### Culture conditions

Synthetic complete (SC) medium contains 0.67% yeast nitrogen base without amino acids, 2% dextrose, 0.1% casamino acids, 50 μg/ml adenine hemisulfate, and either 25 μg/ml uracil (SC-Trp) or 100 μg/ml L-tryptophan (SC-Ura). Supplemented minimal medium (SMM) contains 0.67% yeast nitrogen base without amino acids, 2% dextrose, 20 μg/ml adenine hemisulfate, 20 μg/ml uracil, 20 μg/ml methionine, 30 μg/ml lysine. SMM also contains, depending on selection needs, 20 μg/ml histidine, 100 μg/ml leucine, and/or 20 μg/ml tryptophan, as indicated. SLac medium lacking histidine contains 0.67% yeast nitrogen base without amino acids, 1.2% NaOH, a volume of lactic acid sufficient to subsequently bring the pH to 5.5, 20 μg/ml adenine hemisulfate, 20 μg/ml uracil, 20 μg/ml methionine, 30 μg/ml lysine, 100 μg/ml leucine, and 20 μg/ml tryptophan. Solid media also contain 1.7% bacteriological agar. Cells were incubated at 30°C unless otherwise indicated. For serial dilution assays, strains in logarithmic proliferation phase were diluted to an OD600 of 0.1, and 4 μL of this dilution and three serial five-fold dilutions were spotted to solid medium. Experiments have been carried out at 30°C unless otherwise noted.

### Fis1p TA mutant library construction

Recombination-based cloning (Oldenburg *et al.* 1997) was used to generate constructs expressing Gal4-sfGFP-Fis1p under control of the *FIS1* promoter and mutated at one of 27 positions within the Fis1p TA. Two DNA segments generated by PCR were fused in this recombination reaction. The 5’ portion was amplified by PCR from template plasmid b100 using primer 698 and the appropriate primer (rvsposX) listed in Table S4. The 3’ section was generated from template b100 using primers 517 and the relevant primer (fwdposX) listed in the same table. PCR products were recombined into *Not*I-linearized pKS1 by co-transformation of vector and PCR products into strain MaV203. The sub-library for each Fis1p TA position was generated individually by selection of Trp+ clones in liquid medium, with a portion of each transformation reaction plated to solid SC-Trp medium to confirm recombination and transformation efficiency. To generate the total pool prior to selection for Gal4p-mediated transcription, equal numbers of cells, as determined by OD600 measurement, were taken from overnight cultures of each sub-library and combined within the same liquid culture.

### Deep mutational scanning of the Fis1p TA library

The pool of constructs containing Fis1p TA mutations was cultured for four generations in SC-Trp medium, SC-Ura medium, or SMM-Trp-His medium containing 0 mM, 5 mM, 10 mM, or 20 mM 3-AT. Plasmids present under each culture condition were then recovered from 10 OD600 units of cells. To harvest each plasmid library, cells were pelleted at 4,000 g for 3 min, then washed with 5 ml 0.9 M D-sorbitol and resuspended in 1 ml of 0.9 M D-sorbitol. One “stick-full” of zymolyase 20T (Amsbio, Abingdon, United Kingdom), was added, and cells were incubated at 37°C for 45 min. Cells were again collected at 4,000 g for 3 min and processed using a plasmid purification kit (GeneJET Plasmid Miniprep Kit, Thermo Scientific, Waltham, USA) according to the manufacturer’s instructions. Primers 882 and 883 were used to amplify the genomic region encoding the Fis1p TA from each plasmid pool. Using the provided PCR products, next-generation, paired-end sequencing was performed by Microsynth (Balgach, Switzerland) on a MiSeq Nano (2x150v2). The resulting FASTQ output can be found at [to be uploaded to Dryad Digital Repository and link provided - upload requires manuscript acceptance. Data are also available upon reviewer request]. FASTQ output from paired ends, stripped of adaptor sequences, was combined into a single segment using the PANDAseq assembler version 2.8 (Masella *et al.* 2012). The TRIM function (trimmer Galaxy tool version 0.0.1) was performed using the resources of the Galaxy Project (Goecks *et al.* 2010) in order to remove sequences not directly encoding the defined Fis1p and stop codon. Further processing in Microsoft Excel (Redmond, USA) subsequently allowed conversion of DNA sequence to amino acid sequence and removal of those TAs with more than one amino acid alteration from further analysis. Enrichment values reflect, at a given amino acid position, the ratio of the fraction of amino acid counts following selection to the fraction of amino acid counts in the starting library. Enrichment values are not derived through comparisons across different amino acid positions. Counts for the native amino acid at each position were set as the total number of TA counts for which all amino acids were WT within a given selected pool. When calculating enrichment values, TA amino acid replacements for which there were zero reads in the SC-Trp sample had their value changed to one in order to allow possible detection of enrichment under selective conditions by preventing division by zero. Heat maps were generated using the Matrix2png utility (Pavlidis and Noble 2003).

### Microscopy

For epifluorescence microscopy, cells in the logarithmic phase of culture proliferation were examined using an Eclipse 80i microscope with a 100X Plan Fluor objective and linked to a DS-Qi1Mc camera (Nikon, Tokyo, Japan). Cells were cultured in SMM medium appropriate for plasmid selection. Exposure times were automatically determined, and images were captured using NIS-Elements version AR 3.2. mCherry fusions are driven by the *ADH1* promoter and universally contain Fis1p amino acids 119-128 linking mCherry to the Fis1p TA, the region of Fis1p that is necessary and sufficient for mitochondrial insertion (Mozdy *et al.* 2000; Beilharz 2003; Kemper *et al.* 2008; Förtsch *et al.* 2011) or to alternative TAs. All images of mCherry expression were brightness adjusted in Adobe Photoshop CS5 (Adobe, San Jose, California) to an equivalent extent, except when the mCherry-BAX(TA) signal was assessed. For presentation of data associated with that specific fusion protein, the ‘autolevels’ adjustment was used. Scoring of mitochondrial morphology was performed blind to genotype. To promote Fis1p-dependent mitochondrial fragmentation, sodium azide was added at a concentration of 500 μM for 60 min before fluorescence microscopy (Fekkes *et al.* 2000; Klecker *et al.* 2015).

### Genetic assessment of Fis1p functionality

Cells lose mtDNA and the ability to proliferate on non-fermentable medium when mitochondrial fusion is blocked unless mitochondria division is also abrogated (Sesaki and Jensen 1999; Fekkes *et al.* 2000; Mozdy *et al.* 2000; Tieu and Nunnari 2000). Strain CDD688 (Mutlu *et al.* 2014) harbors chromosomal deletions in *FZO1* and *FIS1* and a CHX-counterselectable plasmid expressing *FZO1*. Upon removal of plasmid-expressed *FZO1*, cells will maintain mtDNA and respire unless functional *FIS1* is also present to allow mitochondrial division. To assess the functionality of Fis1p variants containing TA mutations, strain CDD688 was transformed with plasmids expressing WT *FIS1* or variants mutated within the Fis1p TA. Transformants were cultured overnight in SMM-His medium lacking CHX to permit cells to lose the *FZO1*-encoding plasmid. Serial dilutions were then spotted to SLac-His + 3 μg/mL CHX (“lactate / no fusion”) and incubated for 5 d to test for maintenance of mtDNA following counterselection for *FZO1*, with cell proliferation indicating a lack of Fis1p function. As a control for cell proliferation under conditions not selective for mtDNA maintenance, an equal number of cells was also spotted to SMM-Trp-His medium (“glucose / fusion”) and incubated for 2 d.

## ACKNOWLEDGEMENTS

We thank Gülayse Ince Dunn, Bengisu Seferoglu, Güleycan Bal, Funda Kar, and Sara Nafisi for comments on this manuscript. This work was supported by a European Molecular Biology Organization Installation Grant (2138) to CDD, a European Research Council Starting Grant (637649-RevMito) to CDD, and by Koç University.

## SUPPLEMENTAL FIGURE AND TABLE LEGENDS

**Figure S1.**
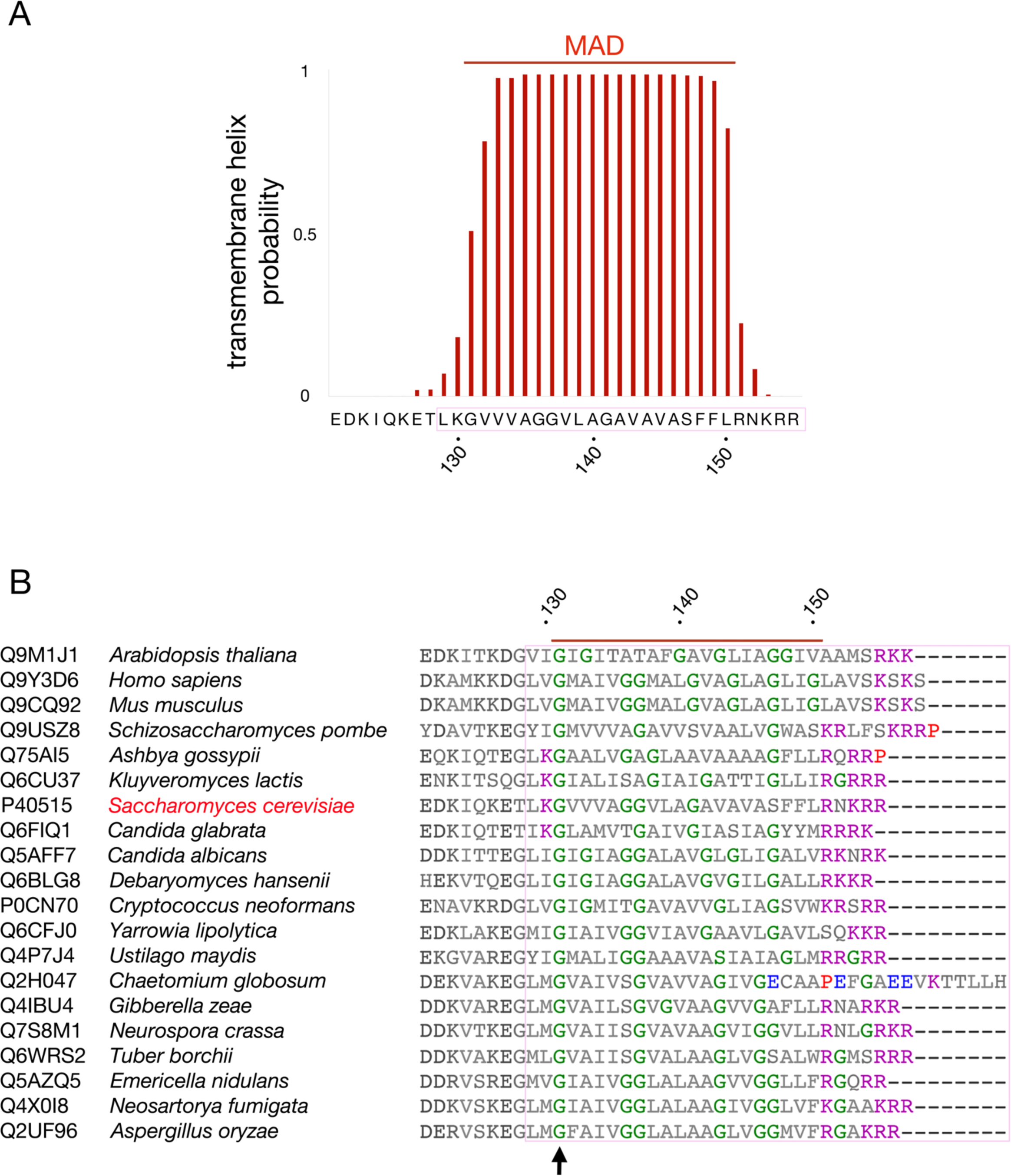
Structural and sequence characteristics of the Fis1p TA. (A) The Fis1p TA is predicted to be a transmembrane helix. The indicated amino acid sequence was tested using the TMHMM Server version 2.0 (Krogh *et al.* 2001), and individual amino acid scores were plotted. The predicted MAD is indicated. (B) Glycines are abundant among Fis1p orthologs. Fis1 proteins from the listed species (Uniprot access numbers listed) were aligned using Clustal Omega (Soding 2005). Glycines are colored green, positively charged amino acids are colored purple, negatively charged amino acids are colored blue, and proteins are colored red. A red bar over the alignment indicates the predicted MAD, and a black arrow denotes a potentially conserved glycine. The region necessary and sufficient for Fis1p insertion is bracketed in pink.

**Figure S2.**
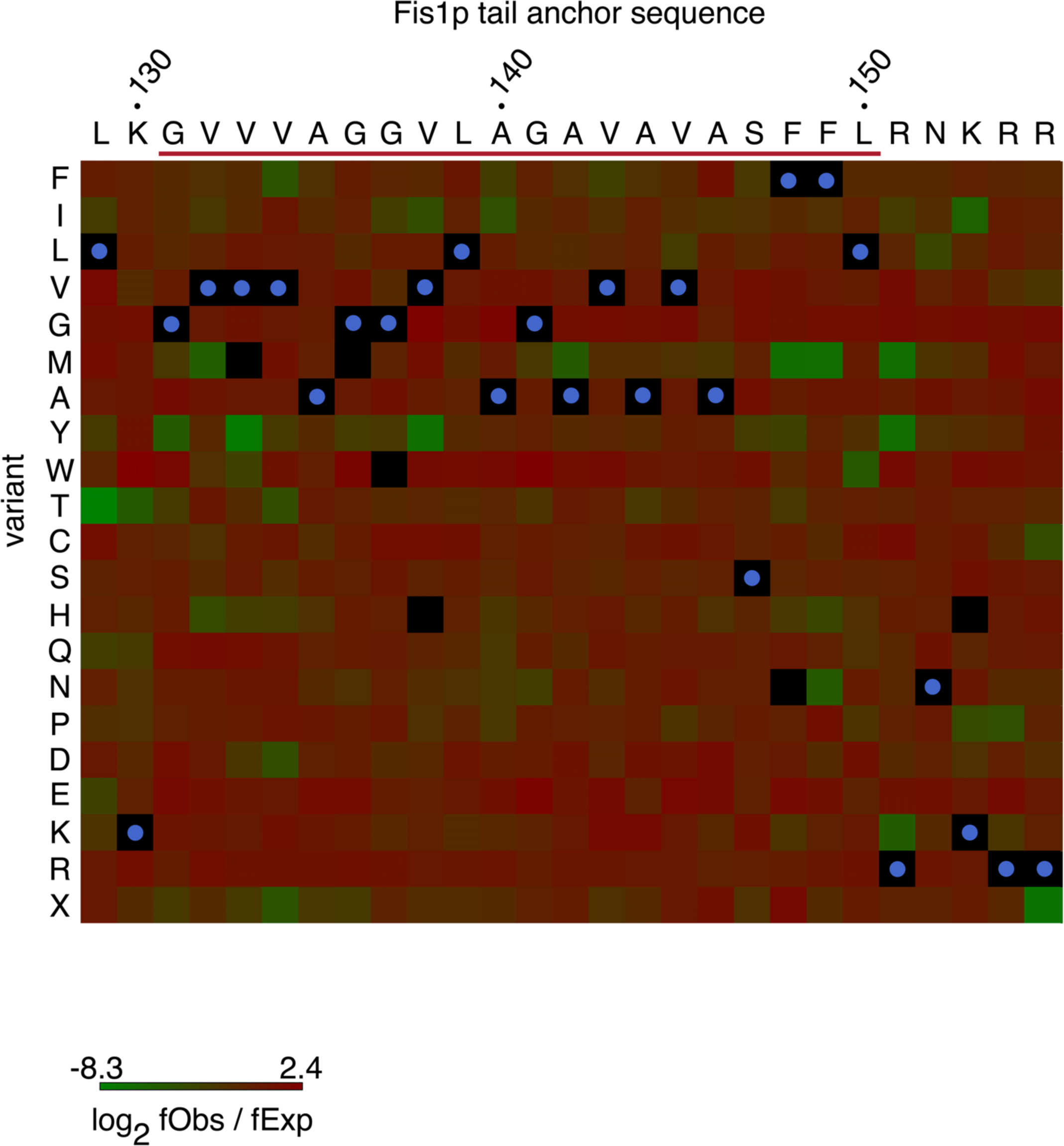
Amino acid replacement representation within the Gal4-sfGFP-Fis1p TA library. The fraction of counts representing each amino acid replacement in the starting SC-Trp library (fObs) was compared to the fraction that would be expected based on randomized codon recovery (fExp). Native amino acids are represented by a black square with a blue dot. Amino acid replacements with no representation in the library are represented by empty black squares. The predicted MAD is indicated by a red line.

**Figure S3.**
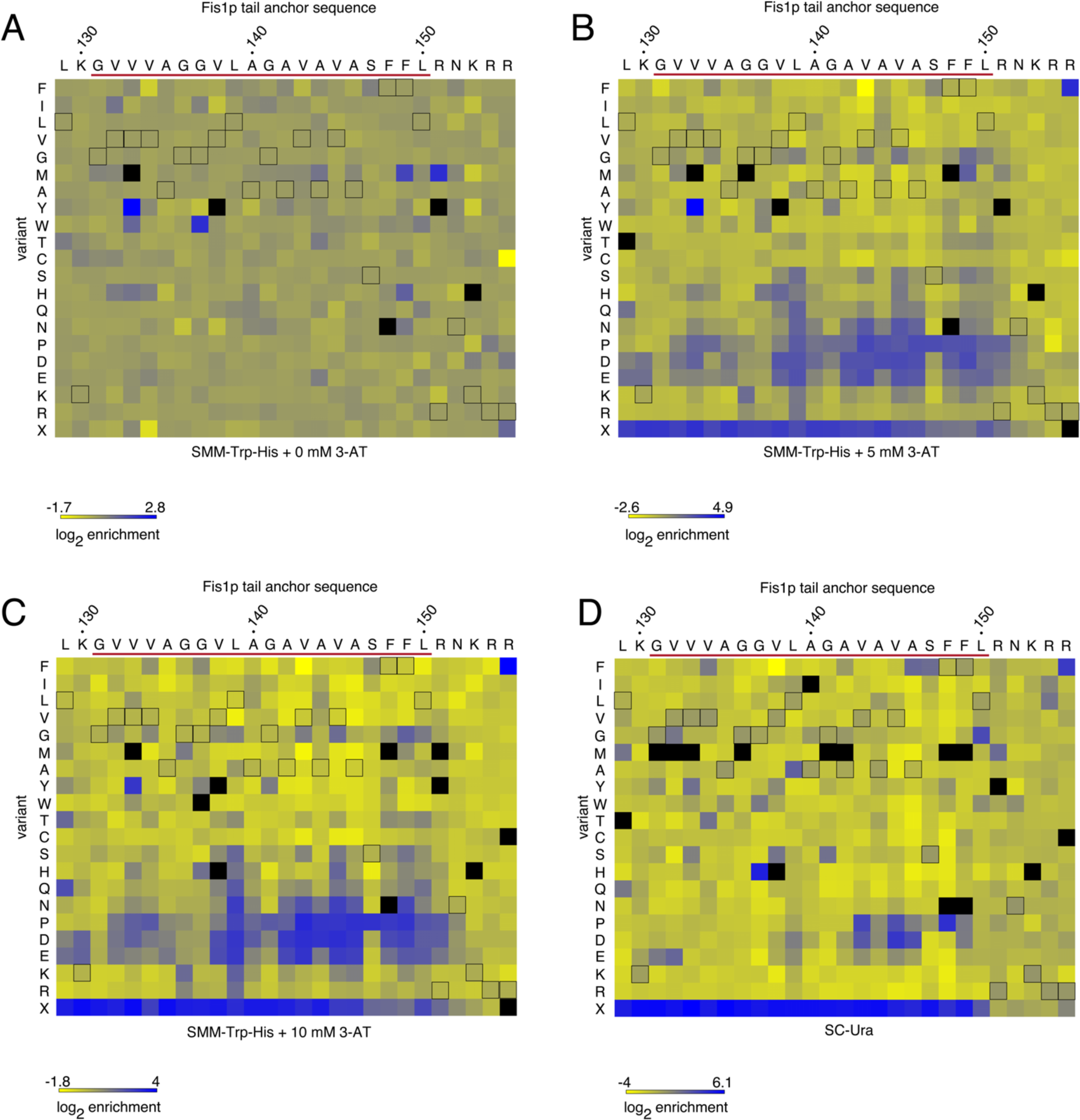
Amino acid replacement frequencies within the Fis1p TA under differing selective pressure. Quantification of replacement in SMM-Trp-His medium without 3-AT (A), containing 5 mM 3-AT (B), containing 10mM 3-AT (C), or in SC-Ura medium (D). In all panels, black outlines indicate the native amino acid at each position within the Fis1p TA. Amino acid replacements not detectable under selective conditions are denoted by black, filled squares. The predicted MAD is indicated by a red line.

**Figure S4.**
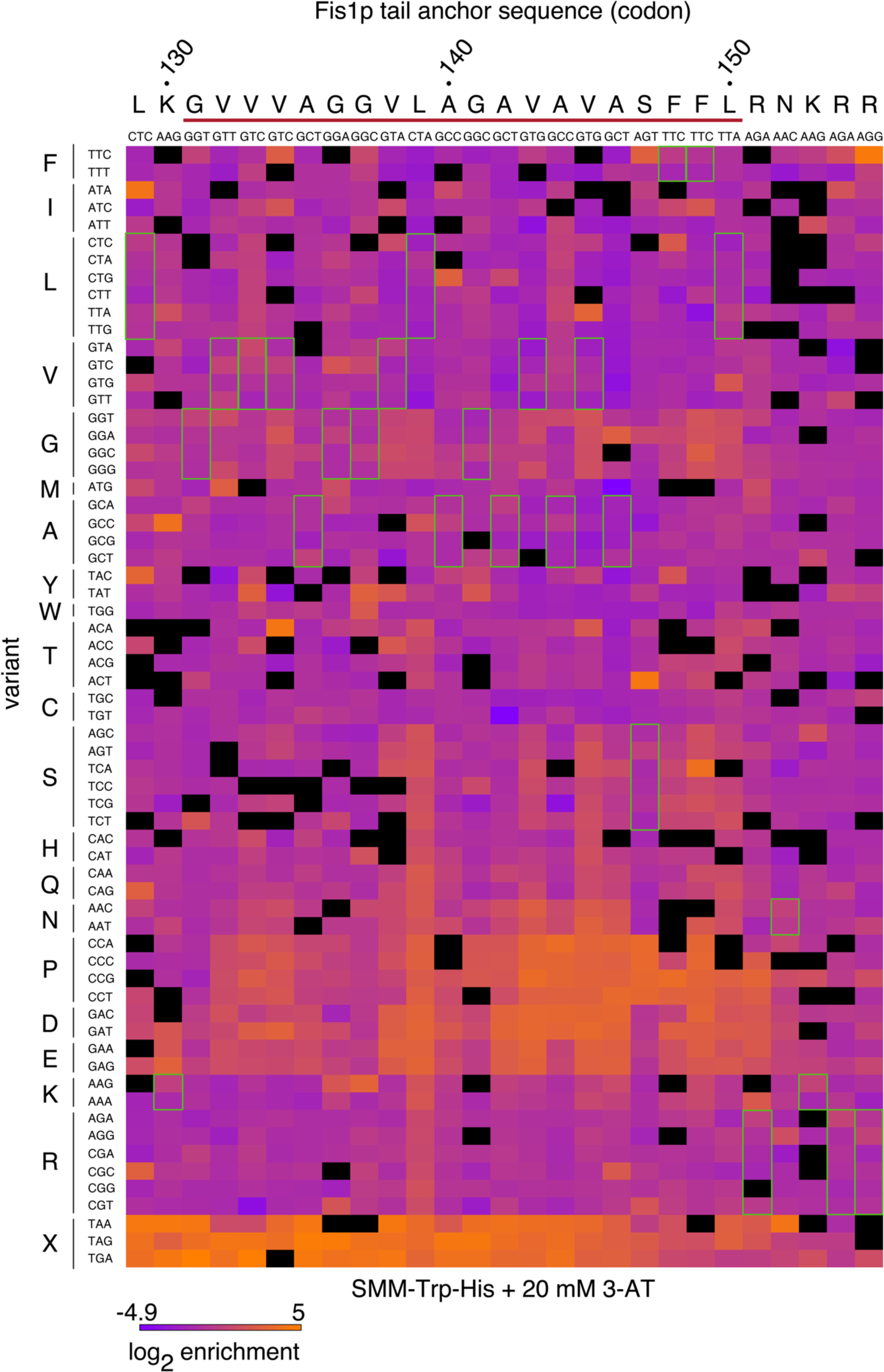
Little evidence exists for codon-level control of Gal4-sfGFP-Fis1p TA targeting. The log2 of enrichment values for each codon following selection of the Fis1p TA library in SMM-Trp-His medium containing 20 mM 3-AT are illustrated. Green outlines denote the native amino acid at each position. Codon replacements with no representation in the library following selection are represented by empty black squares. The predicted MAD is indicated by a red line.

**Figure S5.**
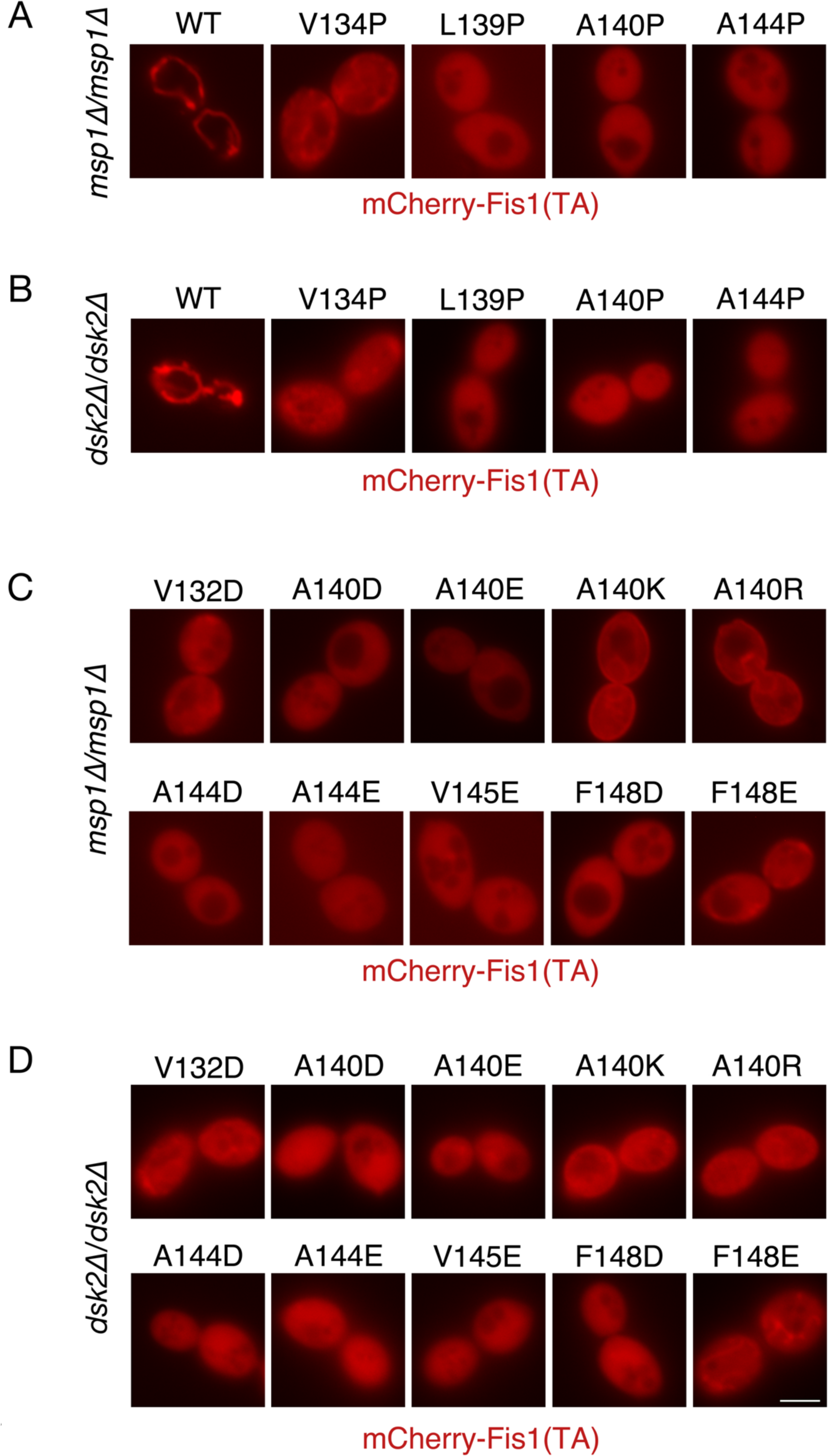
Deletion of Msp1p or Dsk2p does not allow recovery of mitochondrial localization by poorly targeted mCherry-TA variants. (A) Deletion of the Msp1p extractase does not allow tail-anchored proteins mistargeted due to proline substitutions to return to mitochondria. Cells from *msp1Δ/msp1Δ* strain CDD1044 expressing pHS1 (signal not shown) and plasmids b109 (WT), b208 (V134P), b135 (L139P), b210 (A140P), or b211 (A144P) were examined to determine mCherry-TA location. (B) Deletion of ubiquilin ortholog Dsk2p does not allow mislocalized, proline-containing Fis1p TAs to target to mitochondria. Cells from *dsk2Δ/dsk2Δ* strain CDD1179 were transformed and analyzed as in (A). (C) Deletion of Msp1p does not allow mitochondrial localization of mistargeted Fis1p TAs carrying charge replacements. Strain CDD1044, deleted of Msp1p, was transformed with the following plasmids encoding mCherry fused to the mutant TAs: b192 (V132D), b196 (A140D), b197 (A140E), b198 (A140K), b199 (A140R), b200 (A144D), b201 (A144E), b134 (V145E), b204 (F148D), b205 (F148E). Transformants were examined as in (A). (D) Removal of Dsk2p fails to allow mutant TAs containing charge substitutions to re-localized to mitochondria. Strain CDD1179 was transformed with plasmids and analyzed as in (C). Scale bar, 5 μm.

**Figure S6.**
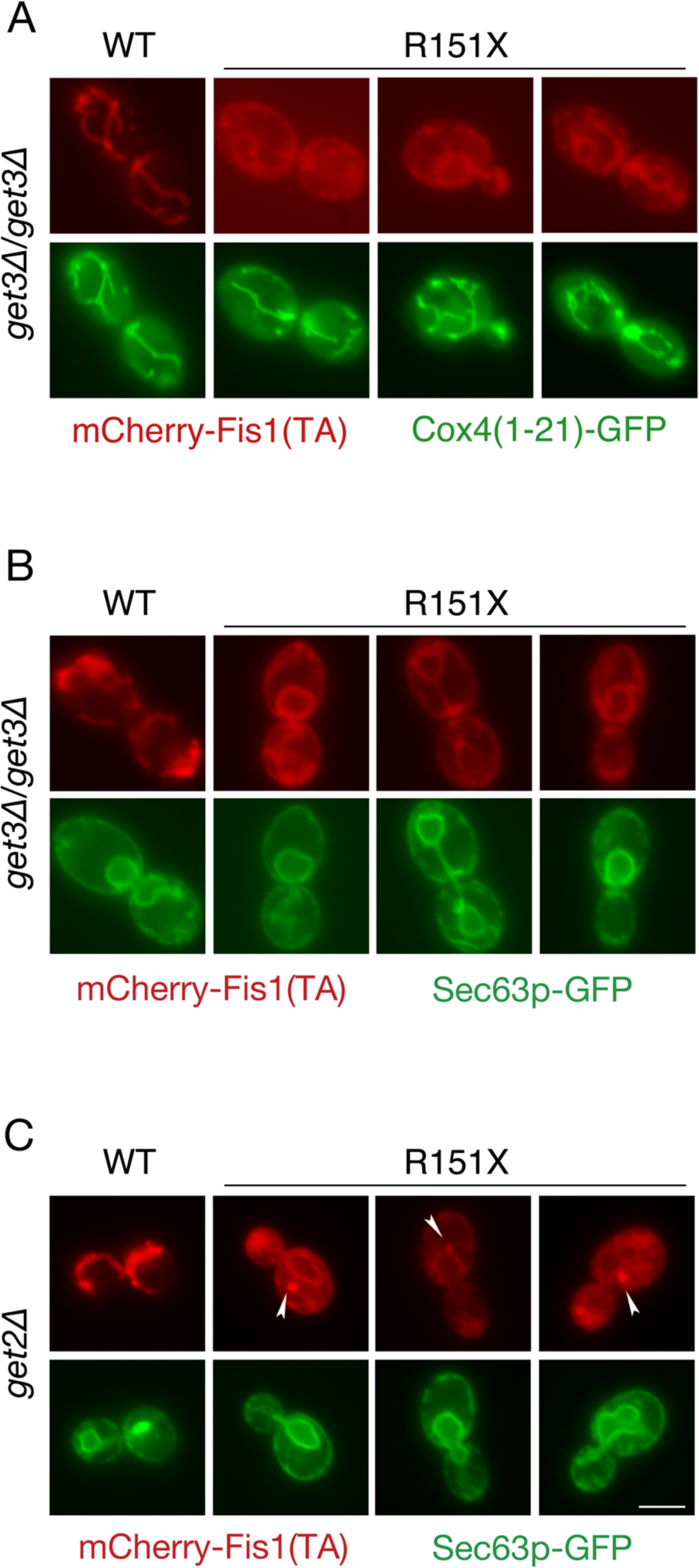
A mutated Fis1p TA lacking the positively charged carboxyl-terminus localizes to the ER independently of Get3p or Get2p. (A) Get3p is not required for ER localization of the R151X TA. *get3Δ/get3Δ* strain CDD1033 expressing mCherry fused to a WT Fis1p TA from plasmid b109 or expressing mCherry fused to a Fis1p TA lacking the positively charged carboxyl-terminus (R151X) from plasmid b254 were examined, and mitochondria were labelled with GFP expressed from pHS1. (B) *get3Δ/get3Δ* strain CDD1033 carrying plasmids b109 or b254 were examined by expression of Sec63p-GFP from plasmid pJK59 as in Figure 6C. (C) Get2p is not necessary for ER localization of the Fis1p TA lacking carboxyl-terminal, positively charged amino acids. *get2Δ* strain CDD948 was transformed with plasmid pJK59 and either b109 or b254, then imaged as in Figure 6C. White arrows indicate puncta containing truncated mCherry-Fis1(TA). Scale bar, 5 μm.

**Figure S7.**
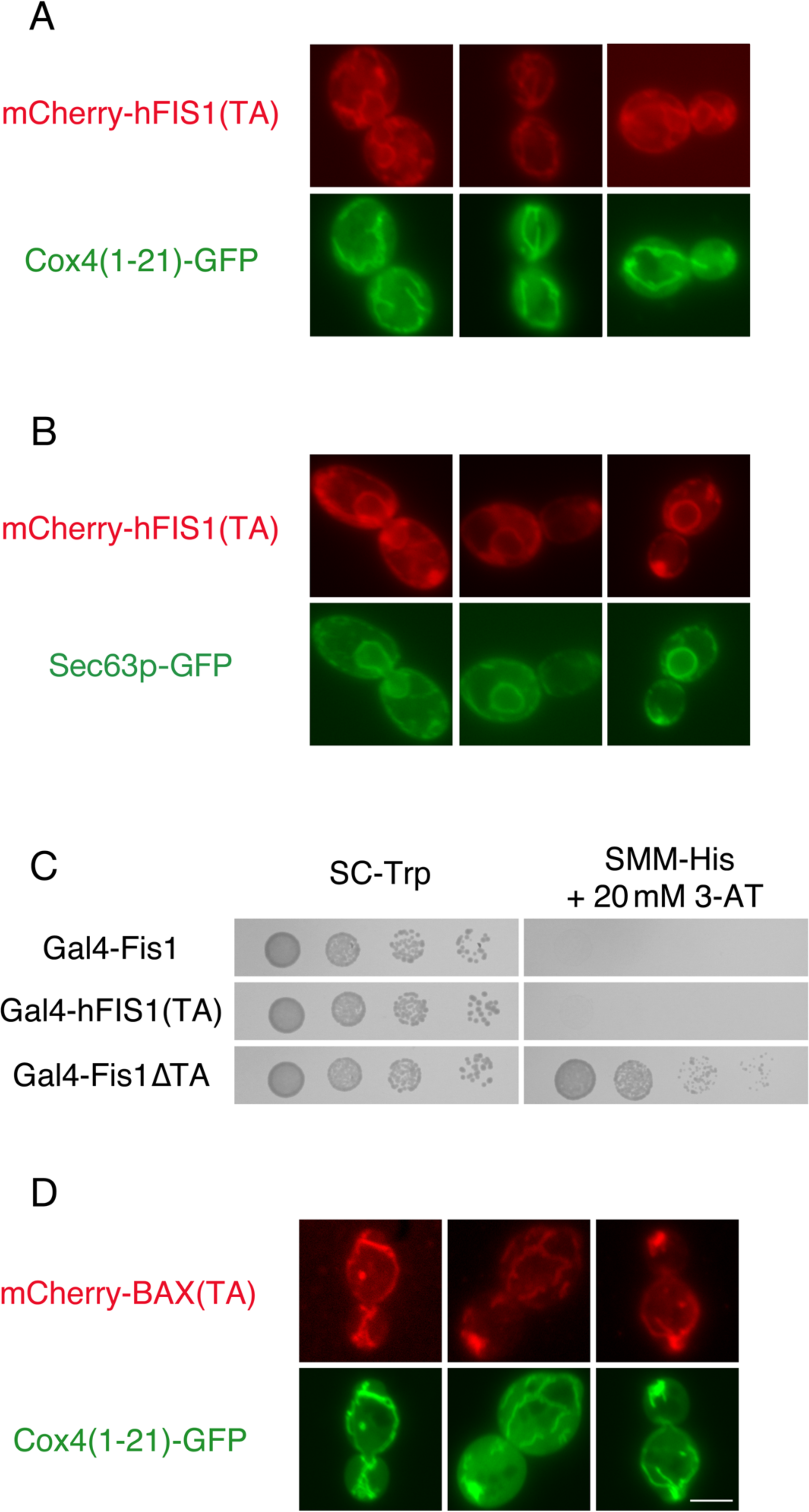
The TAs of human proteins do not universally target to mitochondria in yeast. (A) The hFIS1 TA is mistargeted to the ER in *S. cerevisiae*. Strain CDD961 containing plasmid b257 expressing mCherry fused to the hFIS1 TA was analyzed as in Figure 2B. (B) Strain CDD961 cured of pHS1 and carrying plasmids pJK59 and b257 was imaged as in Figure 6C. (C) The hFIS1TA does not permit activity of a fused Gal4p within the nucleus. Strain MaV203 was transformed with plasmid b100 (Gal4-sfGFP-Fis1), b258 [Gal4-hFIS1(TA)], or plasmid b101 (Gal4-sfGFP-Fis1ΔTA) and assessed as in Figure 5A. (D) The BAX TA can target to mitochondria in *S. cerevisiae*. Strain CDD961 transformed with plasmid b255, which expresses mCherry fused to the BAX TA, was examined as in Figure 2B. Scale bar, 5 μm.

**Figure S8.**
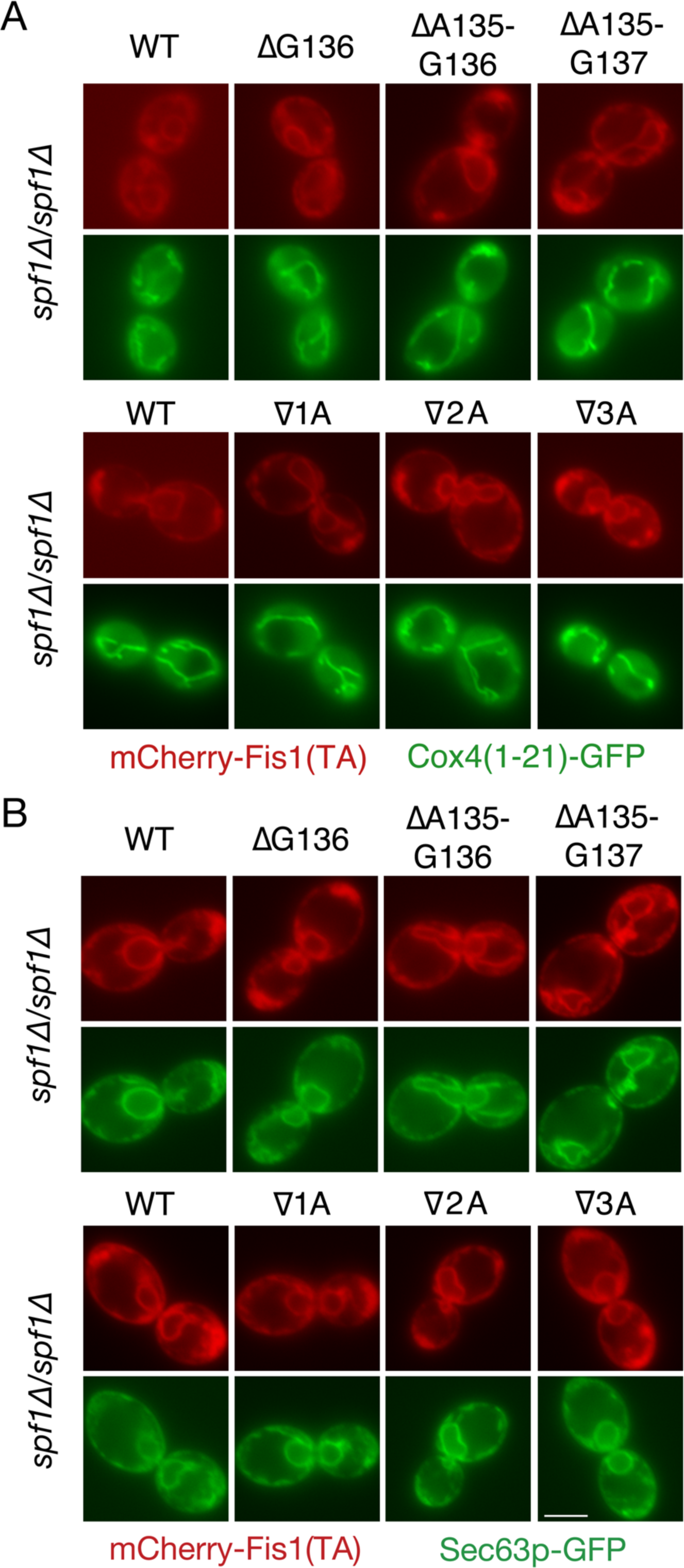
mCherry fused to Fis1p TAs of varying length remains targeted to the ER in cells lacking Spf1p. (A) Cells of *spf1Δ/spf1Δ* strain CDD1031 expressing mCherry-TA fusions from plasmids b109 (WT), b232 (ΔG136), b233 (ΔA135-G136), b234 (ΔA135-G137), b235 (∇1A), b236 (∇2A), or b237 (∇3A), were examined as in Figure 2B. (B) The same mCherry-linked TAs were analyzed in CDD1031 cells expressing Sec63p-GFP from pJK59. Scale bar, 5 μm.

**Figure S9.**
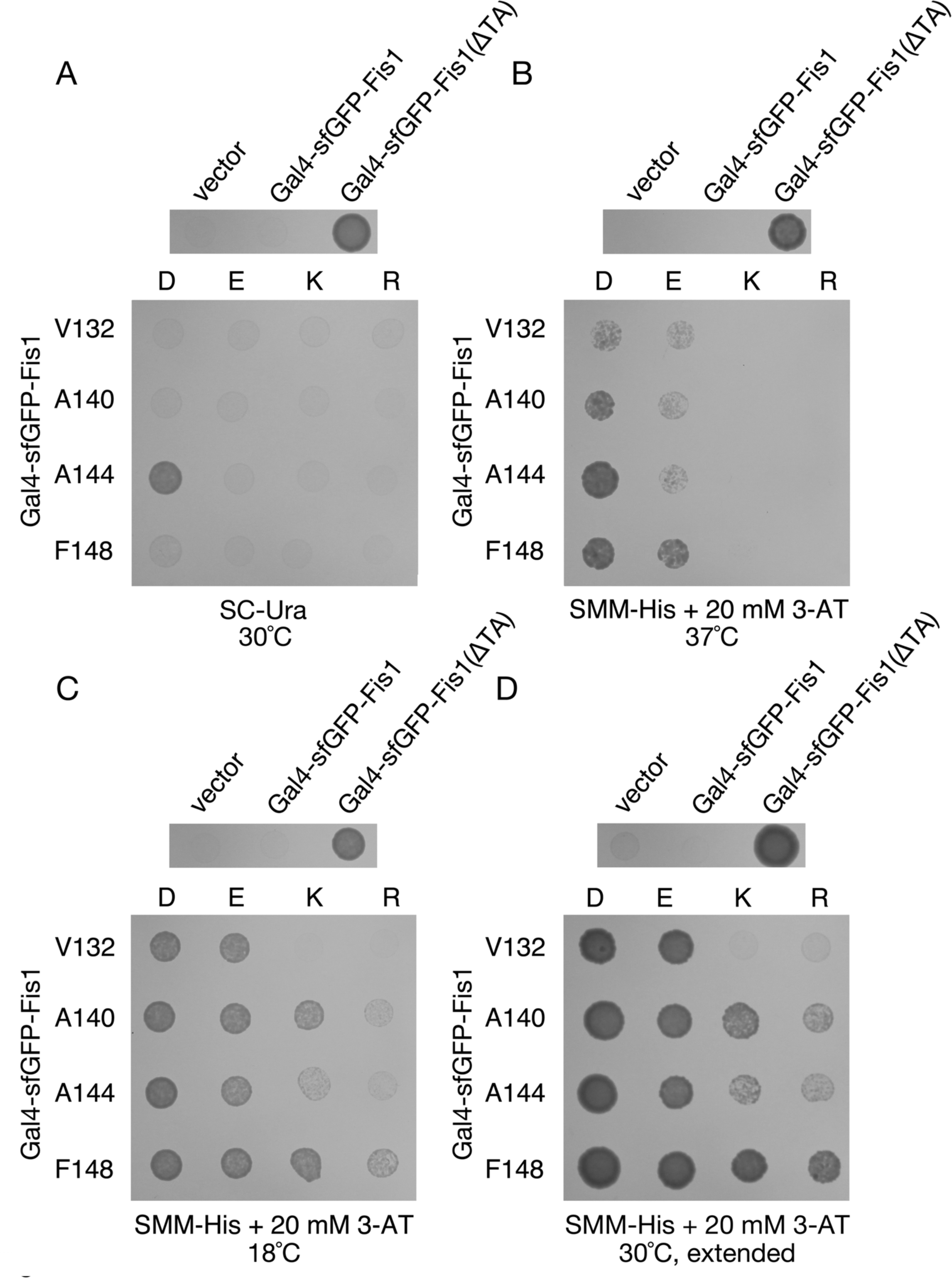
Temperature and selection-dependent proliferation in the presence of charges within the Fis1p MAD. (A) The A144D replacement within Gal4-sfGFP-Fis1p allows for strong Gal4-dependent proliferation under stringent selective conditions. Cells spotted in Figure 8A were also spotted to SC-Ura medium and incubated at 30°C for 2 d. (B) Elevated temperature further distinguishes the behavior of cells expressing Gal4-sfGFP-Fis1p carrying positively charged or negatively charged amino acid replacements. Cells used in Figure 8A were also spotted to SMM-His + 20mM 3-AT and incubated at 37°C for 4 d. (C) Cells were treated as in (B), with incubation at 18°C for 4 d. (D) Extended incubation of the SMM-His + 20 mM 3-AT plate shown in Figure 8A for 4 d at 30°C.

**Figure S10.**
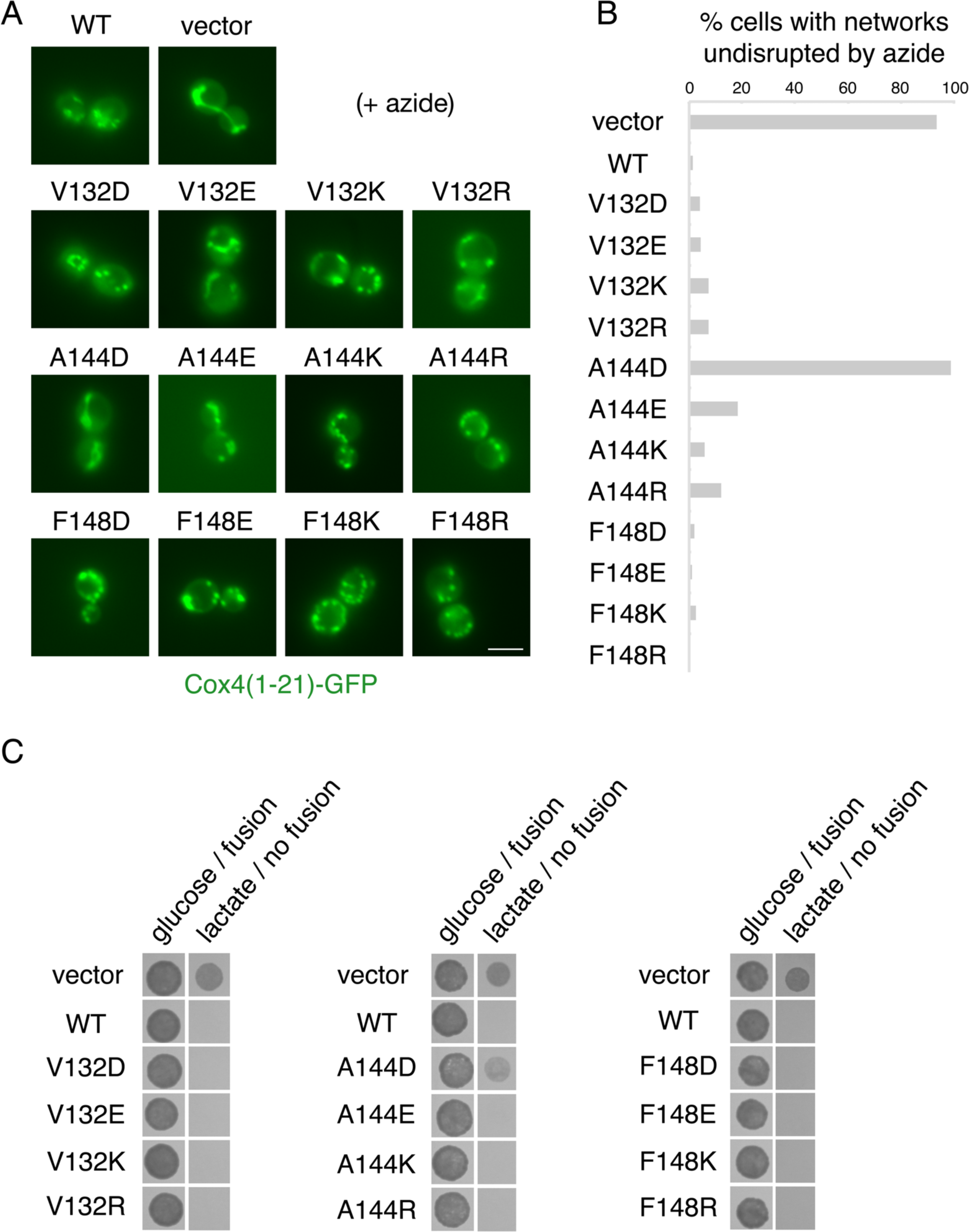
Positive and negative charge replacements within the Fis1p TA can generally support Fis1p function. (A) Normal mitochondrial morphology can be maintained even when charges are placed within the hydrophobic MAD of the Fis1p TA. *fis1Δ* strain CDD741, in which mitochondria were visualized by expression of mitochondria-targeted GFP from plasmid pHS12, was transformed with vector pRS313, a plasmid expressing WT Fis1p (b239), or plasmids expressing Fis1p containing the indicated TA alterations (plasmids b240-251). Cells were treated with sodium azide to provoke Fis1p-dependent mitochondrial fragmentation, and representative images are shown. Scale bar, 5 μm. (B) Cells treated as in (A) were scored for the maintenance of a mitochondrial network (n=200 cells). (C) A genetic assay of Fis1p function demonstrates that charged residues within the TA allow Fis1p activity. *fzo1Δ fis1Δ* strain CDD688, carrying a CHX-counterselectable, *FZO1*-expressing plasmid, was transformed with the *FIS1*-expressing plasmids enumerated above. After allowing cells to lose the *FZO1*-encoding plasmid, serial dilutions were spotted to SLac-His medium containing 3 μg/mL CHX to counterselect against *FZO1* expression (“lactate / no fusion”) and incubated for 5 d to test for maintenance of mtDNA. Cell proliferation indicates a lack of Fis1p activity. To control for cell number spotted, cells from the same culture were placed on SMM-Trp-His medium (“glucose / fusion”) and incubated for 2 d.

**Figure S11.**
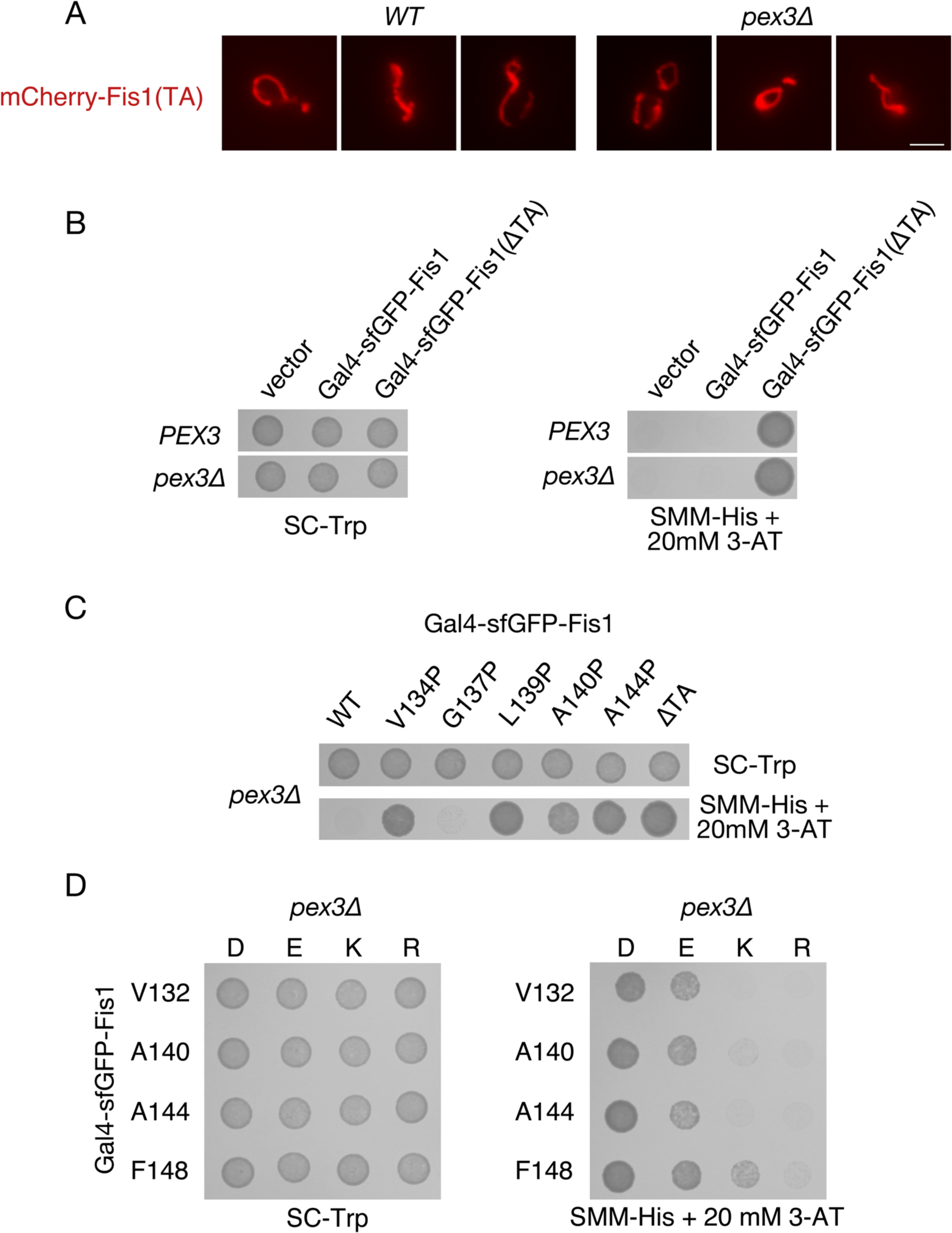
A block of peroxisomal membrane biogenesis does not lead to re-localization of the Fis1p TA to the cytosol or nucleus. (A) *WT* strain BY4742 and *pex3Δ* strain CDD974 were transformed with plasmid b109, expressing mCherry fused to a wild-type Fis1p TA, and imaged by fluorescence microscopy. Scale bar, 5 μm. (B) Deletion of Pex3p does not lead to increased Gal4-sfGFP-Fis1p transcriptional activity. *PEX3* strain MaV203 and isogenic *pex3Δ* strain CDD1172 were transformed with plasmids b100 (Gal4-sfGFP-Fis1), b101 [Gal4-sfGFP-Fis1(ΔTA)], or empty vector pKS1 and treated as in Figure 8A. (C) Relative activity of Gal4-sfGFP-Fis1p containing proline substitutions does not change upon Pex3p removal. Strain CDD1172 expressing Gal4-sfGFP-Fis1p variants from plasmids b100 (WT), b188 (V134P), b189 (G137P), b129 (L139P), b190 (A140P), b296 (A144P), or b101 (ΔTA) was treated as in (B). (D) Relative differences in Gal4-sfGFP-Fis1p transcriptional activity among charge-substituted TAs do not change in the absence of Pex3p. Strain CDD1172 transformed with plasmids encoding the indicated mutations within the Fis1p TA (plasmids b173-b187 and b295) was treated as in (B).

**Table S1.** Tail anchor sequence counts from individual pools.

**Table S2.** Strains used during this study.

**Table S3.** Plasmids used for experiments during this study.

**Table S4.** Oligonucleotides used in this study.

